# Mechanistic implications of the ternary complex structural models for the photoenzyme protochlorophyllide oxidoreductase

**DOI:** 10.1101/2023.09.04.556209

**Authors:** Aoife Taylor, Shaowei Zhang, Linus O. Johannissen, Michiyo Sakuma, Robert S. Phillips, Anthony P. Green, Sam Hay, Derren J. Heyes, Nigel S. Scrutton

## Abstract

The photoenzyme protochlorophyliide oxidoreductase (POR) is an important enzyme for understanding biological H-transfer mechanisms. It uses light to catalyse the reduction of protochlorophyllide (Pchlide) to chlorophyllide, a key step in chlorophyll biosynthesis. Although a wealth of spectroscopic data have provided crucial mechanistic insight about the light-driven reaction chemistry, a structural rationale for POR photocatalysis has proved more challenging and remains hotly debated. Recent structural models of the ternary enzyme-substrate complex, derived from crystal and electron microscopy data, show differences in the orientation of the Pchlide substrate and the architecture of the POR active site that have significant implications for the catalytic mechanism of Pchlide reduction. Here, we have used a combination of computational and experimental approaches to investigate the compatibility of each of these structural models with the hypothesised reaction mechanisms and propose an alternative structural model for the cyanobacterial POR-Pchlide-NADPH ternary complex based on these findings. Through detailed site-directed mutagenesis studies we show that a strictly conserved Tyr residue, which has previously been proposed to act as the proton donor in POR photocatalysis, is not likely to be involved in this step of the reaction but is crucial for Pchlide binding. Instead, an active site Cys residue is important for both hydride and proton transfer reactions in POR and is proposed to act as the proton donor, either directly or through a water-mediated network. Moreover, a conserved Gln residue is found to be important for Pchlide binding and ensuring efficient photochemistry by tuning its electronic properties, likely by interacting with the central Mg atom of the substrate. This optimal ‘binding pose’ for the POR ternary enzyme-substrate complex illustrates how light energy can be harnessed to facilitate enzyme catalysis by this unique enzyme.

## INTRODUCTION

Photoenzymes use light for catalysis but are rare in nature [1]. However, our understanding of how light energy can be harnessed to facilitate enzyme catalysis is now becoming crucial for the design and engineering of non-natural photo-biocatalysts [1, 2]. One such photoenzyme is the light-dependent protochlorophyliide oxidoreductase (POR), which uses coenzyme NADPH to catalyse the reduction of the C17-C18 double bond of protochlorophyllide (Pchlide) to chlorophyllide (Chlide), an important step in the biosynthesis of chlorophyll [3]. POR plays an important role in the formation of the photosynthetic apparatus and the subsequent development of the plant, and is therefore crucial to all life on Earth [3]. In addition, POR is an important enzyme for studying mechanisms of H-transfer in biology [4–11]. Some photosynthetic organisms possess a ‘dark’ Pchlide oxidoreductase that consists of three separate protein subunits and can catalyse the same reaction in the absence of light in an ATP-dependent manner [3].

A mechanistic rationale for the light-driven catalytic reaction of POR is only possible with a detailed understanding of the structure of the POR–NADPH–Pchlide ternary complex. Although these remained elusive for many years, the recent crystal [12, 13] and electron microscopy [14] structures of the enzyme have provided much-needed insight. However, how the Pchlide substrate is orientated in the active site is still unclear [12, 14]. The crystal structures of the *apo* and NADPH-bound forms of cyanobacterial POR enzymes revealed a typical dinucleotide binding Rossmann-fold with a central β-sheet and multiple flexible loops [12, 13]. In the absence of a Pchlide-bound crystal structure, computational modelling was used to propose how Pchlide is likely to bind in the active site [12] (PDB 6RNV). It has been hypothesised that Pchlide is orientated with polar functional groups forming hydrogen bonds with hydrophilic residues located in a deep binding pocket. The hydrophobic region of Pchlide was thought to interact with hydrophobic residues to form a hydrophobic patch on the surface of the protein [12]. This model (**Figure 1A**) was shown to be consistent with previous mutagenesis studies [15, 16] and highlighted how a number of active site residues that are crucial for POR activity (e.g. Tyr193, Lys197 and Thr145) may interact with the propionic acid group at the C17 position of the Pchlide substrate (<4 Å) Pchlide [12].

**Fig. 1.**
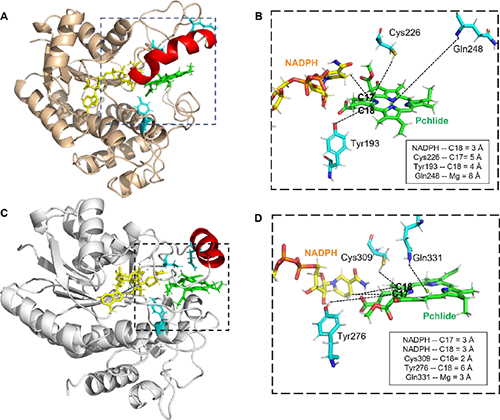
Recent structural models for the POR-Pchlide-NADPH ternary complex for plant and cyanobacterial PORs. **A.** POR from *T. elongatus* crystal structure with docked Pchlide (6RNV) [12]. **B.** Close-up of the active site of POR from 6RNV model showing key residues, Tyr193, Cys226 and Gln248, investigated in the current work together with respective distances to the C17/C18 region and central Mg atom of Pchlide shown by black dashed lines. NADPH is shown in orange, Pchlide in green and active site residues in cyan. **C.** POR from *A. thaliana* based on a cryo-EM model (7JK9) [14]. **D.** Analogous residues compared to those in panel **B** from 7JK9 model together with respective distances to the C17/C18 region and central Mg atom of Pchlide shown by black dashed lines.

A subsequent cryo-EM structure of POR from the plant *Arabidopsis thaliana* (PDB 7JK9) has shown how Pchlide-bound POR oligomers form helical filaments with the lipid bilayer [14]. This helical assembly appears to be responsible for the characteristic curved architecture of the prolamellar bodies in photosynthetic membranes [14] and is consistent with energy transfer between neighbouring Pchlide molecules, which might contribute to increased reaction efficiency [17, 18]. However, structural differences exist between the cryo-EM structure and the previous structural model that may have significant implications for the POR catalytic mechanism. One difference is that two flexible regions of the POR protein which close over the hydrophobic edge of Pchlide to form a ‘lid’ [19], are in a more ‘closed’ conformation in the cryo-EM derived ternary complex of POR [14]. These ‘lid’ regions position the Pchlide in a geometry that is optimal for photocatalysis and their movement is proposed to trigger larger conformational change that facilitate POR oligomer formation [19]. Moreover, the structural models also differ significantly in the conformation of the bound Pchlide molecule in the active site [12, 14]. The strictly conserved residues, Tyr276 and Lys280 (numbering now according to *A. thaliana* POR), remain in close proximity to the active site, although the Lys residue is too far away (>4 Å) from the propionic acid group to H-bond in this model. Conversely, the conserved, catalytically important Cys309 residue appears optimally positioned to participate in the reaction chemistry as it is located < 5 Å from the C18 position of the Pchlide substrate.

The differences between the models are likely to have significant implications for the current proposed hypotheses for the mechanism of catalysis by POR. Previous spectroscopic studies have led to a proposed catalytic mechanism whereby Pchlide photochemistry triggers hydride transfer from NADPH to the C17 position of Pchlide on a timescale of ∼ 500 ns [6, 7, 16]. This is followed by proton transfer to the C18 position on the microsecond timescale (∼ 50 μs), either directly from the putative active site Tyr193 proton donor or via active site water molecules [5–7, 15]. The roles of several active site residues in this mechanism have also been supported by previous mutagenesis studies [16]. However, the publication of the new structural model (PDB 7JK9) for the plant POR-Pchlide-NADPH ternary complex has led to an alternative proposed catalytic mechanism [14]. In this case, it is suggested that the hydride from NADPH could attack either the C17 or C18 position of Pchlide as the pro-*S* face of the nicotinamide ring is found on a horizontal plane with the C17-C18 double bond of Pchlide [14]. Due to the known stereochemistry of chlorophyll [20–22], if the NADPH attacks from above the Pchlide plane in this model then it must be to the C18 position and if it attacks from below the plane it must be to the C17 position. The subsequent proton is then proposed to be derived from either Cys309 or Ser228 [14]. Ser228 is not conserved across all POR enzymes, and is instead, a threonine (Thr145, numbering in *Thermosynechococcus elongatus* POR) in cyanobacterial PORs (**Figure S1**), which is unlikely to act as a proton donor. However, previous mutations to this Thr residue have resulted in inactive POR enzymes and has previously been shown to participate as part of a proton relay mechanism in other closely related enzymes [16]. Cys309 is fully conserved throughout all POR enzymes and interestingly, when the equivalent residue in *T. elongatus* POR, Cys226, was previously mutated to a Ser residue it appeared to influence the hydride and proton transfer chemistry [23]. To complicate matters further, a recent computational study suggested yet another alternative mechanism [11]. In this scenario, Pchlide photochemistry leads to electron transfer from the conserved Tyr residue to the C18 position of Pchlide, followed by a subsequent proton transfer, also from the Tyr residue, then electron and hydride transfer from NADPH [11].

In the present work we have explored the veracity of each of the structural models using a combination of site-directed mutagenesis, non-natural amino acids, steady-state, cryogenic trapping and laser photoexcitation experiments. These have been augmented with further computational methods, such as docking, DFT and MD simulations. The role of the catalytic Tyr residue as the putative proton donor [15] has been explored in more detail, with evidence suggesting it is unlikely to perform this role in POR. In addition, an in-depth analysis has been carried out of two other active site residues, Cys226, which is potentially important for hydride and proton transfer, and Gln248, a residue that hasn’t previously been investigated but is located close to (and may interact with) the central Mg atom of Pchlide in the cryo-EM structural model of the POR ternary enzyme-substrate complex. Based on these findings, an alternative structural model for the cyanobacterial POR-Pchlide-NADPH ternary complex is presented, together with a discussion of the implications for the potential mechanism of Pchlide reduction.

## RESULTS AND DISCUSSION

### Computational analysis of alternative structural models for the POR-Pchlide-NADPH ternary complex

Molecular dynamics (MD) simulations were used to investigate the two POR ternary enzyme-substrate complex structural models [12, 14] and to assess the associated implications for the reaction mechanism of the enzyme. Both models are structurally stable across the simulations, as the root mean square distribution (RMSD) of Pchlide in the active site was less than 3 Å (**Figure S2**). Analysis of the MD simulations for the cyanobacterial POR structural model (PDB 6RNV) showed that the putative catalytic residues, Tyr193, Thr145 and Lys193 [15, 16], were in close proximity to the propionic acid group at the C17 position of Pchlide and remained at a plausible distance (<4 Å) for H-bonding interactions over the course of the simulation (**Figure 2A and B**). These interactions are proposed to be important to position the Pchlide substrate in the correct orientation for the reduction reaction to occur [12]. The alignment of Pchlide, NADPH and Tyr193 was also analysed over the MD simulations to investigate whether the distance and bond angle for hydride and proton (using Tyr193 as putative proton donor in this case) transfer were suitable for reaction chemistry (**Figure 2C**). The distance between the C4 *pro-S* hydrogen of NADPH and the C17 position of Pchlide fluctuates between 2-3 Å with an angle of 140-160° (**Figure 2D**), both of which are compatible with the efficient transfer of a hydride. For proton transfer the distance between the putative donor, Tyr193, and the C18 position of Pchlide was slightly longer, between 3-4 Å, but the angle of transfer was mainly found to be below 100°, which is significantly lower than the optimal transfer angle of approximately 180° [24]. In the recent structure of the plant POR, Cys309 and Ser228, which correspond to Cys226 and Thr145 in the cyanobacterial POR, were postulated to be potential proton donors [14]. In the cyanobacterial structural model, the proton transfer distance from Thr145 to C18 (preferred position as it attacks from below the ring) was shorter (3-4 Å) than the equivalent distance from Cys226 to C17 (preferred position as it attacks from above the ring, 5-7 Å) (**Figure 2E**). Conversely, the angle of proton transfer was more optimal for Cys226 to C17 (most frequent ∼160°) than for Tyr193 to C18 (**Figure 2F**). However, it is possible that minor conformational changes following the hydride transfer step may optimise the proton transfer chemistry in this structural model.

**Fig. 2.**
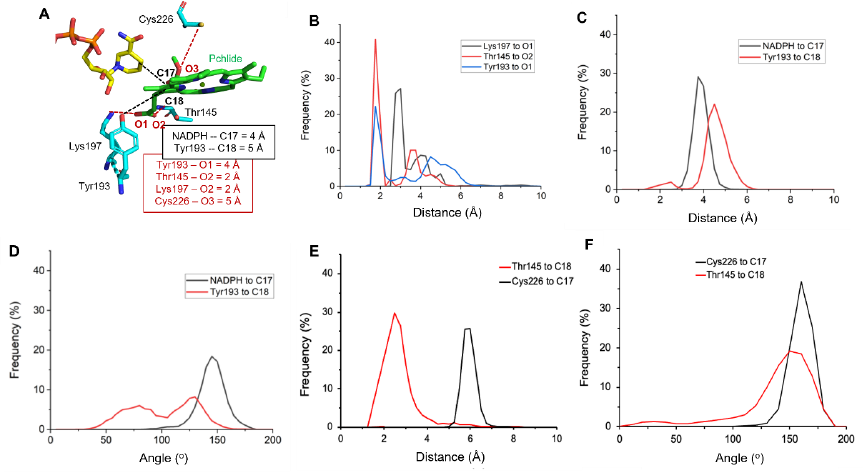
Computational analysis of POR-Pchlide-NADPH ternary complex model from *T. elongatus* (6RNV). **A.** Snapshot from *T. elongatus* POR structural model highlighting key transfer distances (black dashed lines) and potential hydrogen bonds (red dashed lines). **B.** Distances of potential hydrogen bonds from key residues to the propionic group on Pchlide; Lys197 (black), Thr145 (red), Tyr193 (blue). **C.** Frequency polygon of hydride transfer distance from NADPH coenzyme to the C17 position of Pchlide (black) and proton transfer distance from Tyr193 to the C18 position (red). The distance of Lys197 to the Pchlide O3 is 95% > 6 Å and is therefore not included. **D**. Frequency polygon of hydride transfer angle from NADPH coenzyme to the C17 position of Pchlide (black) and proton transfer angle from Tyr193 to the C18 position (red). **E.** Frequency polygon of transfer distance from Cys226 to the C17 position of Pchlide (black) and from Thr145 to the C18 position (red). **F.** Frequency polygon of transfer angle from Cys226 to the C17 position of Pchlide (black) and from Thr145 to the C18 position (red). All MD simulations were run for 10 ns of protein-restrained equilibration and 110 ns unrestrained.

MD simulations on the cryo-EM model of the plant POR-Pchlide-NADPH ternary complex (PDB 7JK9) [14] showed that the putative catalytic residues, Tyr276 and Ser228 were in close proximity to the propionic acid group at the C17 position of Pchlide and remained at an optimal distance (<4 Å) for H-bonding interactions over the course of the simulation, whereas Lys279 was in closer proximity to the NADPH coenzyme than Pchlide. (**Figure 3A and B**). The hydride transfer distance from the *pro-S* hydrogen at the C4 position of NADPH to both C18 and C17 carbons of the Pchlide substrate was too far for efficient transfer to occur (>6 Å) (**Figure 3C**), whilst the hydride transfer angle was also sub-optimal (<120°) (**Figure 3D**). Although this implies that the model is not fully optimised or is in a non-reactive conformation, these distances / angles may change slightly upon photoexcitation of the Pchlide molecule. The proton transfer distance was measured from two potential proton donors, Cys309 and Ser228, which were previously highlighted from the cryo-EM model [14]. These residues are positioned above and below the Pchlide plane and would therefore, only be able to transfer to the C18 or C17 position of Pchlide, respectively to form Chlide with the correct stereochemistry [25]. A wider distribution of distances was observed for Ser143 to the C17 of Pchlide (3-9 Å) compared to Cys309 to C18 (5-8 Å), although neither demonstrated a favourable transfer distance of <4 Å (**Figure 3E**). The angle for proton transfer was highly unlikely from Cys309 to C18 (broad range of angles from 0-180°) but possible for Ser228 to C17 (most frequently 160° which is close to the optimum angle of 180°) (**Figure 3F**). Proton transfer from the other putative proton donor, Tyr276, to either the C17 or C18 positions of Pchlide is highly unlikely in this conformation due to the large transfer distances of >5 Å (**Figure 3G**), despite an optimal angle of attack (most frequently 160°) (**Figure 3H**). Hence, in this structural model it is likely that the proton would be transferred via a water network that is potentially coordinated by the active site residues, Ser228, Tyr276 and Cys309.

**Fig. 3.**
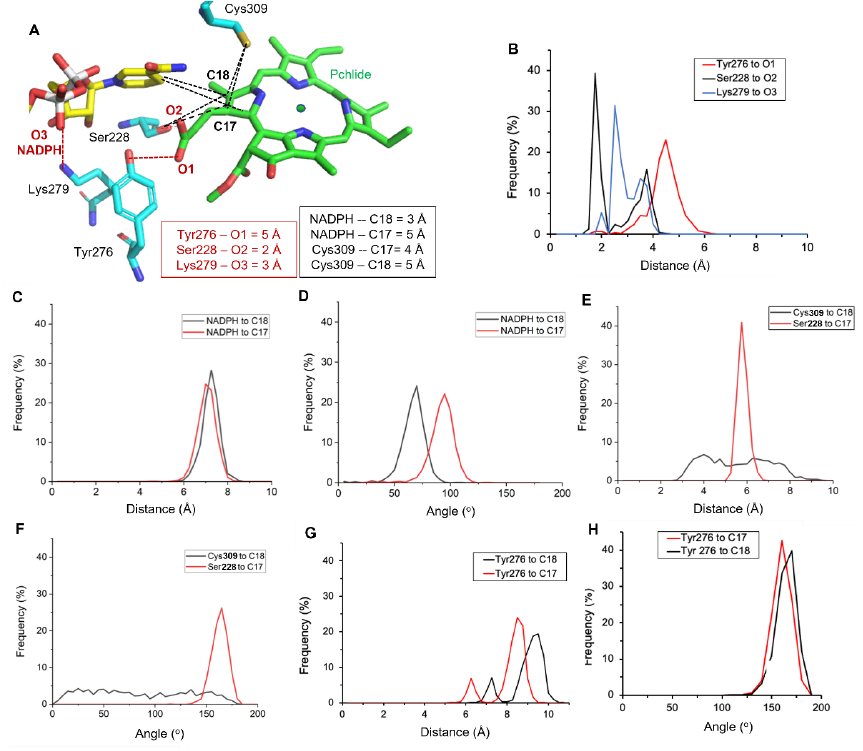
Computational analysis of POR-Pchlide-NADPH ternary complex model from *A. thaliana* (7JK9). **A.** Snapshot from *A. thaliana* POR structural model highlighting key transfer distances (black dashed lines) and potential hydrogen bonds (red dashed lines). **B.** Frequency polygon of potential hydrogen bonding distances from Tyr276 to O1 of the Pchlide propionic acid group (black), Ser228 to O2 of Pchlide propionic acid group (red) and Lys279 to O3 of NADPH (blue). The distance of Lys279 to the Pchlide propionic acid group is 95% >6 Å and is therefore not included. **C.** Frequency polygon of hydride transfer distance from NADPH to either C17 (red) or C18 (black) positions of Pchlide. **D.** Frequency polygon of hydride transfer angle from NADPH to either C17 (red) or C18 (black) positions of Pchlide. **E.** Frequency polygon of hydrogen transfer distance from Cys309 to C18 position of Pchlide (black) or from Ser228 to C17 position (red). **F.** Frequency polygon of hydrogen transfer angle from Cys309 to C18 position of Pchlide (black) or from Ser228 to C17 position (red). **G.** Frequency polygon of proton transfer distance from Tyr276 to either C18 (black) or C17 (red) positions of Pchlide. **H.** Frequency polygon of proton transfer angle from Tyr276 to either C18 (red) or C17 (black) positions of Pchlide. All MD simulations were run for 10 ns of protein-restrained equilibration and 110 ns unrestrained.

As the two structural models were both of reasonable stability in MD simulations (converging on <3 Å for Pchlide RMSD) this implies that multiple binding conformations of Pchlide might be possible for POR enzymes. To determine whether it was possible for Pchlide to adopt more than one binding pose, the substrate was docked into the POR crystal structure from *T. elongatus* [12]. The top three docked conformations of Pchlide based on the best calculated fit were selected, along with a Pchlide binding pose based on the cryo-EM structural model [14] (**Figure S3**). Pose 1 is the conformation published by Zhang et al in their computational simulations [12] and has the highest docking score. Pose 2 is of a similar conformation (propionic acid group in a different plane to the methoxy group) but with a horizontal axis flip of 180° and a rotation of 90°. Pose 3 and Pose 4, on the other hand, have the propionic acid group on the same plane as the methoxy group but are situated differently in the active site (90° horizontal rotation between the two poses). The computed Pchlide binding energy was compared in each case using the relative difference between the energies obtained for each pose across three separate MD simulations (**Figure S4**). Although significant differences between runs resulted in large errors that prevented the determination of a preferred binding pose the data does suggest that it is energetically possible for Pchlide to adopt multiple binding poses in the POR active site. This implies that there is a significant amount of movement in the active site that perhaps, allows the Pchlide to bind in multiple positions, only one of which is a ‘reactive pose’. This hypothesis would be compatible with previous studies that showed that multiple, slow conformational changes occur upon Pchlide binding, which are necessary to position the substrate in an optimal geometry for reaction chemistry and are rate-limiting for overall photocatalysis by POR [26].

### Tyr193 is key Pchlide binding residue but unlikely to act as proton donor in POR catalysis

The active site residue Tyr193 has previously been shown to be crucial for POR activity and has been proposed to play multiple roles in catalysis, including Pchlide binding, photochemistry and as the putative proton donor to the C18 position of Pchlide [8, 15]. In the original structural model [12] Tyr193 was positioned reasonably close (<5 Å) to the C18 position of Pchlide (**Figure 1A and B**), but in the cryo-EM structural model the Pchlide is in a flipped position which positions Tyr193 closer to the C17 carbon with a less favourable proton transfer distance of >5 Å [14] (**Figure 1C and D**). Previous site-directed mutagenesis studies have only focussed on changing this Tyr residue to a Phe or an Ala residue, which are no longer capable of H-bonding or proton transfer [8, 15]. Here, we initially sought to insert natural amino acid residues that may still be able to act as a proton donor, namely Lys, His and Glu, in place of Tyr193. However, steady-state activity measurements showed very low or negligible activity of these variants and Pchlide binding titrations yielded *K*_d_ values of >100 µM in each case (**Figure S5**), which is >20-fold higher than for wild-type [15, 16, 27]. This suggests that the binding of Pchlide in the POR active site is significantly affected by these mutations. Laser photoexcitation measurements showed a major decrease in quantum efficiency for each of the Tyr193 variants (**Figure 4A-C**), meaning it was not possible to measure the rates of hydride and proton transfer for Y193K and Y193E. However, the Y193H variant did provide a sufficient signal and showed a decreased rate of proton transfer (∼0.95 x10^4^ s^-1^) compared to the WT enzyme (∼2.92 x10^4^ s^-1^) (**Figure 4D**), rather than an expected increase for a lower p*K*_a_ residue if Tyr193 acted as the proton donor.

**Fig. 4.**
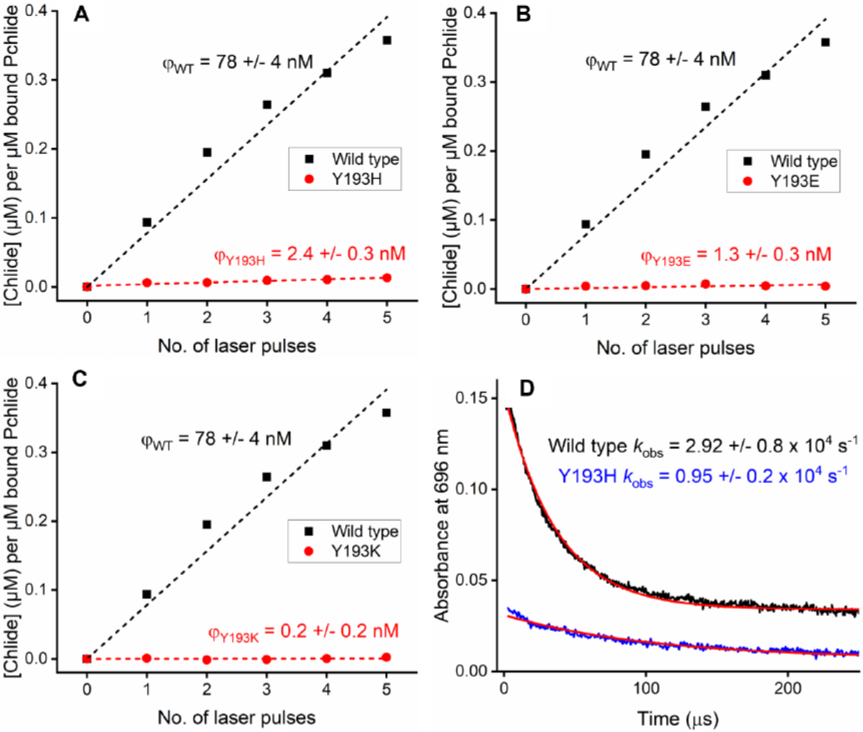
Potential role of Tyr193 in POR photochemistry. **A-C**. Comparison of the relative quantum yield of the wild-type POR-catalysed reaction with Y193H (**A**), Y193E (**B**) and Y193K (**C**) variants from single-turnover laser photoexcitation measurements. The Chlide concentration after each laser pulse was plotted (normalised using bound Pchlide concentration) and fitted to a straight line with the slope as an estimate of the relative quantum yield for the POR reaction. Reaction conditions: 100 µM POR, 15 µM Pchlide, 0.1 % Triton, 1 µM 2-mercaptoethanoland 250 µM NADPH in buffer (150 mm NaCl, 20 mm HEPES, pH 7.5). **D.** Laser flash photolysis transients of wild type (black trace) and Y193H (blue trace) POR showing decrease in absorbance at 696 nm as a measure of the rate of proton transfer [6]. Samples contained 100 µM POR, 15 µM Pchlide, 0.1 % Triton, 1 µM 2-mercaptoethanoland 250 µM NADPH. The *k*_obs_ value was calculated from the average of 5 kinetic transients by fitting to a single exponential.

Due to the significantly impaired binding of Pchlide to the Tyr193 variants, in both the current and previous studies [15], a more subtle approach was sought by exploring the use of non-natural fluorinated Tyr residues (**Figure 5A**) in place of the natural Tyr (p*K*_a_ = 10.0). Fluorinated tyrosine (mono-and tri-fluorinated) was incorporated in a site-specific manner at the Y193 position by using the recently evolved *M. jannaschii* Y-tRNA synthetase/tRNA pair [28] and confirmed by mass spectrometry. Despite low levels of protein expression, a significant reduction in catalytic activity was observed as the level of fluorination of Tyr193 increased, with the tri-fluorinated Tyr193 POR only exhibiting ∼3% of the steady-state activity of the wild-type enzyme at pH 7.5 (**Figure 5B**). The activity of this variant increased at pH 6 compared to pH 7.5, suggesting that the protonation state of Tyr193 may be important for POR catalysis. Single turnover laser flash photolysis measurements were used to provide an estimate for the relative quantum yield for the fluorinated Tyr193 variants, but the tri-fluorinated enzyme had too little activity to measure any product formation in these experiments. Although fluorination of Tyr193 has a significant effect on Pchlide binding, as determined by a lower level of bound Pchlide for mono-fluorinated Tyr193 POR (**Figure S6A**), the concentration of Chlide formed from a single laser pulse does not appear to be affected (**Figure S6B**), implying that there is no effect on the quantum yield for this variant. Taken together, this suggests that it is the Pchlide binding step that is affected in the mono-fluorinated Tyr193 variant, and once bound the reduction reaction chemistry can proceed efficiently. Time-resolved laser spectroscopy on the mono-fluorinated Tyr193 variant showed a proton transfer rate of 2.94 x 10^4^ ± 0.06 s^-1^ (**Figure 5C**), which is almost identical to that of wild type (2.92 x 10^4^ ± 0.08 s^-1^, **Figure 5B**), again suggesting that Tyr193 is not involved in this step. Overall, the data here confirm that Tyr193 is crucial for Pchlide binding but is unlikely to be the proton donor as previously hypothesised [8,12,15], and a different residue, or a water molecule, is likely to play this role in the POR active site.

**Fig 5.**
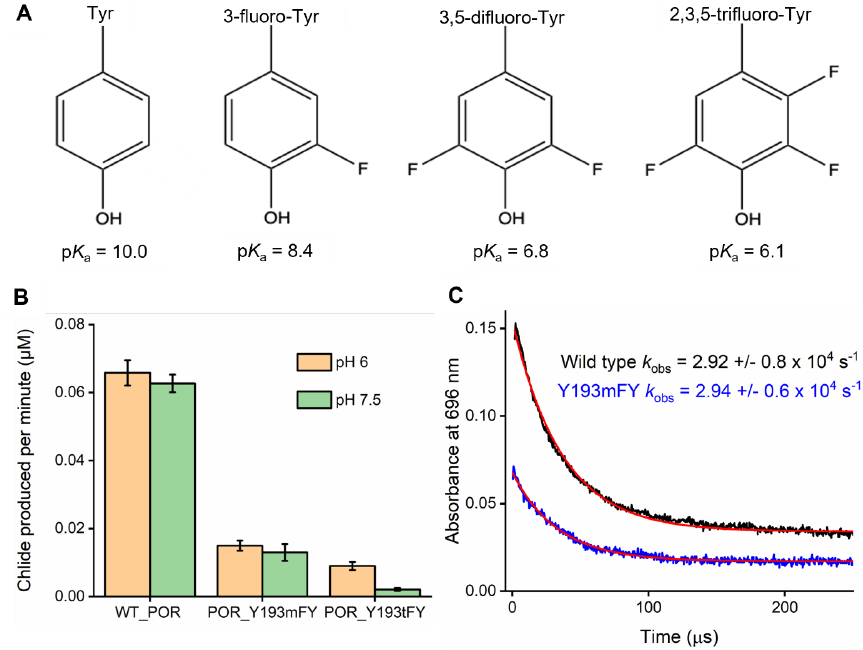
Probing the role of Tyr193 in POR catalysis by using fluorinated Tyr analogues. **A**. Fluorinated Tyr analogues that can be incorporated into proteins using an engineered *M. jannaschii* Tyr-tRNA/aaRS pair [28]. **B.** Steady-state activity of wild-type POR compared to mono-and tri-fluorinated POR variants at pH 6 (orange) and pH 7.5 (green) for samples containing 0.1 µM enzyme, 20 µM Pchlide, 0.1 % Triton, 1 µM 2-mercaptoethanoland 100 µM NADPH in buffer (50 mM MES, 25 mM Tris, 25 mM ethanolamine, 100 mM NaCl). **C.** Kinetic transients showing the decrease in absorbance at 696 nm, which corresponds to the proton transfer step [6], for wild type (black trace) and mono-fluorinated POR variant (blue trace) after laser photoexcitation at 450 nm. Samples contained 100 µM wild type POR or 200 µM mono-fluorinated POR variant, 20 µM Pchlide, 0.1 % Triton, 1 µM 2-mercaptoethanoland 250 µM NADPH in buffer (50 mM MES, 25 mM Tris, 25 mM ethanolamine, 100 mM NaCl). The *k*_obs_ value was calculated from the average of 5 kinetic transients by fitting to a single exponential.

### Cys226 as putative proton donor in POR catalysis?

The active site residue Cys226 has previously been shown to play a key role in the catalytic mechanism of POR [9, 14, 23] and has recently been proposed to act as the potential proton donor in POR catalysis [14]. In a C226S variant of POR, hydride transfer (i.e. to form Pchlide H^-^) gave rise to a different intermediate species, with a broad absorbance peak centred at 530 nm rather than the typical 696 nm absorbing species observed for the wild-type enzyme [9]. It was suggested that this change in absorbance could be due to the Pchlide substrate binding in an alternative conformation (180° flip), thus causing the hydride to transfer to the other carbon (C18 rather than C17) [9]. However, in the original Zhang et al structural model [12] this residue appears to be located too far away from the C17-C18 double bond of Pchlide to play a significant role in the reaction. Conversely, in the recent cryo-EM structure, the equivalent Cys226 residue is located <5 Å from the C18 position and may also interact with other active site residues (equivalent to Tyr223 and Gln248 in *T. elongatus* POR) [14]. As a Ser residue is theoretically still capable of transferring a proton, mutation to an alanine has now been carried out to assess the potential role of Cys226 as a proton donor.

Initial steady-state measurements showed that the C226A variant followed Michaelis-Menten kinetics with a comparable *K*_m_ for Pchlide (∼4.8 μM) to that obtained for the wild-type enzyme (∼4.3 μM), indicating that Cys226 is not a key Pchlide binding residue (**Figure S7**). However, the *k*_cat_ of 0.11 min^-1^ is significantly lower than the wild type activity (*k*_cat_ = 0.18 min^-1^), which is likely to be a result of a lower quantum yield for the C226A enzyme (**Figure S8**). In order to investigate any changes to the catalytic mechanism in the C226A variant the reaction of the C226A-Pchlide-NADPH ternary complex was analysed by cryo-trapping measurements (**Figure 6** and **Figure S9**). Samples were illuminated for 10 mins with a 450 nm LED at a range of temperatures <180 K, which allowed the identification of an initial light-dependent step. The absorbance band at 642 nm, attributed to bound Pchlide [4, 26], disappears and is accompanied by the appearance of a broad absorbance band between 530-580 nm (**Figure 6A**), which is the same intermediate species previously observed for the C226S variant upon hydride transfer [9]. This species is gradually converted to the Chlide product at ∼672 nm in a single light-independent or ‘dark’ step at temperatures ≥ 200 K (**Figure 6B**).

**Fig. 6.**
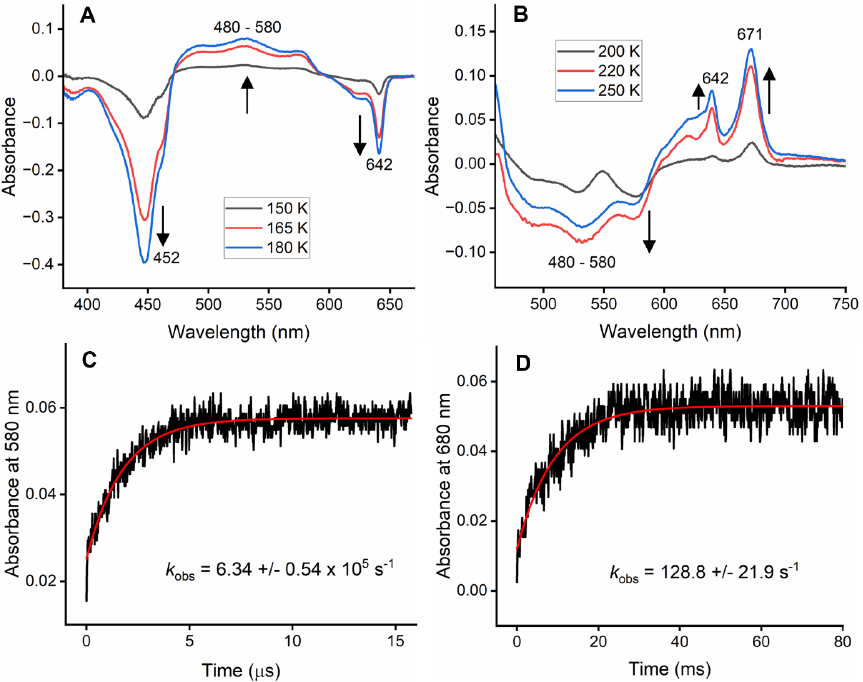
Investigating the role of Cys226 in POR catalysis. **A and B**. Cryogenic absorbance measurements of *T. elongatus* POR C226A variant for samples containing 100 µM C226A, 20 µM Pchlide, 0.1 % Triton, 1 µM 2-mercaptoethanoland 250 µM NADPH in 20 % sucrose, 44 % glycerol, 150 mm NaCl and 20 mm HEPES pH 7.5.**A.** The difference absorbance spectra of the C226A-Pchlide-NADPH ternary complex at 77 K after illumination for 10 min at various temperatures using a non-illuminated sample as a blank. **B.** The difference absorbance spectra of the C226A-Pchlide-NADPH ternary complex at 77 K after illumination for 10 min at 180 K followed by incubation in the dark at various temperatures using the 180 K illuminated sample as a blank. **C** and **D.** Laser flash photolysis transients showing the rates of hydride transfer (increase at 580 nm, **C**) and proton transfer (increase at 680 nm, **D**) for the C226A POR variant. The transients show changes in absorbance after excitation with a laser pulse at 450 nm (15 mJ) for C226A ternary complex samples containing 100 µM C226A, 10 µM Pchlide, 0.1 % Triton, 1 µM 2-mercaptoethanoland 250 µM NADPH. The *k*_obs_ values were calculated from the average of 5 kinetic transients by fitting to a single exponential.

In order to probe the kinetics of these catalytic steps in the C226A variant laser photoexcitation measurements were carried out. The rate of increase at 580 nm was used to measure the hydride transfer step due to the minimal absorbance from the Pchlide triplet state at this wavelength [8, 16] and was calculated to be ∼6.3 x 10^5^ s^-1^ (**Figure 6C**). Measurements repeated in deuterated buffer showed a negligible solvent isotope effect, whereas a kinetic isotope effect (KIE) of ∼ 1.89 ± 0.12 was observed with the deuterated form of the coenzyme, NADPD (**Figure S10**, **Table 1**), confirming that this step corresponds to hydride transfer. The rate of hydride transfer in the C226A variant is approximately 3-fold slower than the rate of hydride transfer in wild-type POR to form the typical intermediate state absorbing at 696 nm (∼2.02 x 10^6^ s^-1^) [6, 7, 9, 15, 16] and comparable to the corresponding rate of formation of the 580 nm absorbing species in the C226S variant (∼0.8 x 10^6^ s^-1^) [9]. The rate of the subsequent proton transfer step, determined by measuring the increase in absorbance at 680 nm associated with the Chlide product [6, 7, 9, 15, 16], was calculated to be ∼125 s^-1^ (**Figure 6D**). The solvent isotope effect and KIE measured for this step in the C226A variant were minimal (**Figure S11**, **Table 1**), implying that the rate may be limited by factors other than proton transfer (e.g. conformational changes). The rate of proton transfer for the C226A variant is >200 fold slower than for wild-type POR (2.92 x 10^4^ s^-1^, **Figure 4D**) [6, 7, 9, 15, 16], and would appear to point towards Cys226 acting as the likely proton donor in POR photocatalysis. Interestingly, the rate of proton transfer was previously found to increase by ∼4-fold for the C226S variant [9, 23]. Alternatively, water may act as the proton donor, in which case alterations to the Cys226 residue may influence the water hydrogen-bonding network, the orientation of the Pchlide in the active site or by changes to any conformational motions required during proton transfer. Either way, it is clear that the Cys226 residue plays a significant role in both hydride and proton transfer and is key for active site chemistry in POR.

**Table 1.** The rates of the hydride transfer and proton transfer steps for the C226A POR variant in the presence of either *pro-S* NADPH or *S*-NADPD and in either H_2_O or D_2_O buffers. Reaction conditions: 100 µM C226A, 10 µM Pchlide, and 250 µM NADPH/D. *The wild-type hydride and proton transfer rates are obtained from [9].

| Sample | Hydride transfer rate (s<sup>-1</sup> x10<sup>5</sup>) |  | Rate of proton transfer (s<sup>-1</sup>) |  |
| --- | --- | --- | --- | --- |
|  | C226A | Wild type* | C226A | Wild type* |
| NADPH in H<sub>2</sub>O | 6.34 ± 0.54 | 20.2 ± 2.1 | 128.8 ± 21.9 | 25,300 ± 1500 |
| NADPD in H<sub>2</sub>O | 3.36 ± 0.30 | 10.0 ± 1.4 | 113.6 ± 22.3 | 25,600 ± 3100 |
| NADPH in D<sub>2</sub>O | 5.78 ± 0.72 | 17.9 ± 2.2 | 96.2 ± 13.3 | 11,500 ± 1100 |
| NADPD in D<sub>2</sub>O | 2.59 ± 2.6 | 8.9 ± 1.2 | 125.8 ± 21.4 | 10,600 ± 400 |

### A key role for Gln248 in POR catalysis

Although the previous data is inconclusive about which structural model is the most feasible for the POR ternary complex, a comparison of the two structures shows that there are key differences in the positioning of Gln248 relative to the Pchlide molecule (**Figure 1**). In the cryo-EM structural model the equivalent Gln248 residue (i.e. Gln331) can be found facing inwards to the active site, surrounded by key ‘lid’ region residues, where it is positioned close enough (<3 Å) to the central Mg atom of Pchlide to potentially form an adduct [14]. Conversely, in the structural model based on the crystal structure, Gln248 is located outside of the active site facing outwards and away from the key ‘lid’ region residues at a distance of 17 Å from the central Mg atom of Pchlide [12]. Hence, site-directed mutagenesis of this residue to produce the Q248A variant would be highly informative about the veracity of each model. Initial steady-state measurements showed a significant reduction in the *k*_cat_ value, and a much higher *K*_m_ value (**Figure S12**). Although this points towards a decrease in the affinity for the Pchlide substrate in the Q248A variant an accurate determination of the *K*_d_ was not possible as the spectral change associated with ternary complex formation is much less pronounced than for the wild-type enzyme. The Pchlide absorbance spectrum does not fully red-shift to 642 nm upon binding, which implies that the electronic properties of the bound substrate are different between the Q248A and wild-type enzymes (**Figure 7A**). Moreover, it was necessary to incubate samples for a much longer period of time (∼1 hr) to observe any spectral change, suggesting that the rate of Pchlide binding, previously shown to be rate-limiting in the overall catalytic cycle [26], is likely to be impaired in the Q248A variant. It is tempting to speculate that these differences may potentially be caused by the loss of interaction to the central Mg atom.

**Fig. 7.**
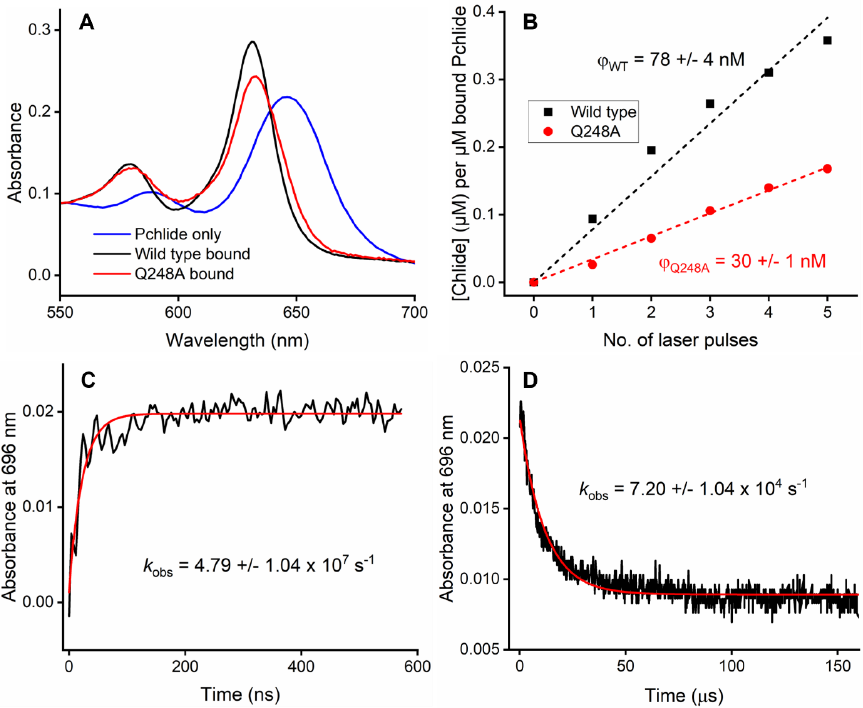
Investigating the role of Gln248 in POR catalysis. **A.** Absorbance spectra of unbound Pchlide (black, λ_max_ = 630 nm), Pchlide bound to Q248A variant (red, λ_max_ = 632 nm) and Pchlide bound to wild type POR (blue, λ_max_ = 642 nm). **B.** Comparison of the relative quantum yield of the wild-type POR-catalysed reaction with the Q248A variant from single-turnover laser photoexcitation measurements. The concentration of Chlide produced per laser pulse was measured (normalised for bound Pchlide concentration) for samples containing 100 µM enzyme, 15 µM Pchlide, 0.1 % Triton, 1 µM 2-mercaptoethanoland 250 µM NADPH. **C and D.** Laser flash photolysis transients showing the rates of hydride transfer (increase at 696 nm, **C**) and proton transfer (decrease at 696 nm, **D**) for the Q248A POR variant. The transients show changes in absorbance at 696 nm after excitation with a laser pulse at 450 nm (15 mJ) for Q248A ternary complex samples containing 100 µM Q248A, 10 µM Pchlide, 0.1 % Triton, 1 µM 2-mercaptoethanoland 200 µM NADPH. The *k*_obs_ values were calculated from the average of 5 kinetic transients by fitting to a single exponential.

Single turnover laser flash photolysis measurements indicate that the relative quantum yield for photocatalysis in the Q248A variant is significantly lower (∼26 %) than the wild-type enzyme (**Figure 7B**). This suggests that Gln248 is essential to facilitate efficient photochemistry. However, despite the reduction in photochemical efficiency, time-resolved spectroscopy was used to measure kinetic transients at 696 nm and showed an ∼20-fold increase in the rate of the hydride transfer step for Q248A (∼4.8 x 10^7^ s^-1^, **Figure 7C**) compared to the wild-type enzyme (∼2.02 x 10^6^ s^-1^) [6, 7, 9, 15, 16]. Moreover, there is also a modest increase in the rate of the proton transfer step (∼6.5 x 10^4^ s^-1^, **Figure 7D**) compared to the wild-type enzyme (∼2.9 x 10^4^, **Figure 4D**). Although it is difficult to fully rationalise these findings at present, it would appear to imply that any changes to the electronic properties of the bound Pchlide substrate caused by mutating the Gln248 residue are likely to have significant impact on the quantum efficiency and rate(s) of the photochemical steps in POR catalysis. Moreover, for Gln248 to influence photochemistry in such a way it suggests that it likely to be positioned close to the Pchlide molecule in the active site and would favour the cryo-EM structural model where it could potentially interact with the central Mg atom of Pchlide [14].

### A new structural model for the T. elongatus POR-Pchlide-NADPH ternary complex and implications for photocatalytic mechanism

Taken together, these results imply that the more ‘closed’ cryo-EM structural model of the ternary enzyme-substrate complex of POR [14] is the most compatible structure for reaction chemistry. The structure of the *T. elongatus* POR ternary enzyme-substrate complex has now been modelled using the plant cryo-EM structure (PDB: 7JK9) [14] as a template (**Figure 8A**). MD simulations have been used to gain an understanding of the implications for the POR catalytic mechanism and to rationalise the experimental findings described above. The RMSD of Pchlide in the active site over 120 ns showed a similar stability to the previous structural models (< 3 Å) (**Figure S13A**). The NADPH coenzyme is situated above the Pchlide plane at a similar, non-optimal distance (>4 Å) and transfer angle (<120°) to both the C17 and C18 of Pchlide (**Figure 8B-E**). Although the hydride from NADPH has previously been shown to transfer to the C17 position [29] this is highly unlikely in the new model as it would lead to the formation of the Chlide product with the incorrect stereochemistry [25] when transferred from above the Pchlide plane. The previously proposed proton donor, Tyr193 [8, 12, 15], appears to be located too far away (∼6 Å) and at a non-optimal transfer angle (∼150°) from the C17 position of Pchlide to perform this role (**Figure 8B and C**), which is in agreement with the earlier experimental findings. Instead, the Tyr193 residue is located in close proximity to the propionic acid sidechain of Pchlide in the new structural model where it is likely to form key H-bonding interactions, thus supporting its crucial role in Pchlide binding (**Figure 8A**). The alternative proposed proton donor, Cys226 [14], is positioned in close proximity (∼2-3 Å) to the C18 position of Pchlide and at a transfer angle (>150°) that is compatible with proton transfer. However, in this case the hydride and proton would both appear to be transferred to the C18 position of the Pchlide substrate. Therefore, unless there are structural rearrangements upon excitation or following hydride transfer it is difficult to rationalise how this active site structure is compatible with reaction chemistry to produce Chlide with the correct stereochemistry.

**Fig. 8.**
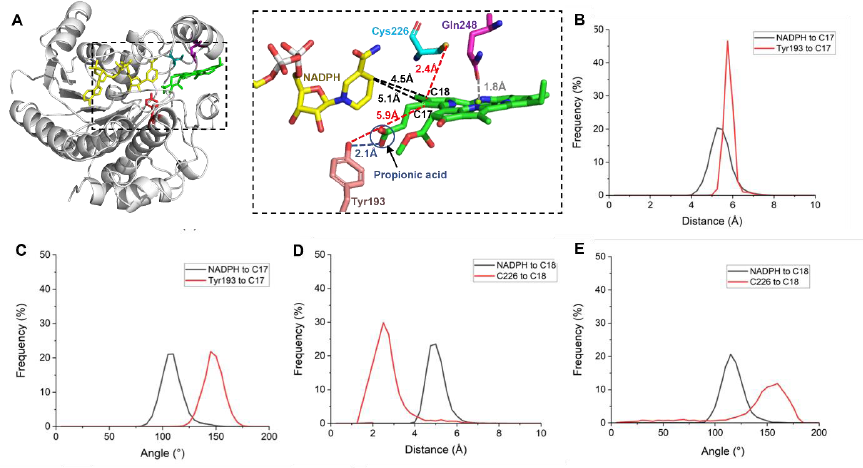
New structural model of POR-Pchlide-NADPH ternary complex from *T. elongatus*. **A.** Overall structure (left) of *T. elongatus* POR with Pchlide (green), NADPH (yellow) and the key residues, Tyr 193 (red), Cys226 (blue), Gln248 (purple), shown. A close-up of active site (right) shows key distances of NADPH and residues from Pchlide. **B.** Frequency polygon of H-transfer distance from Tyr193 (black) or NADPH (red) to the C17 position of Pchlide. **C.** Frequency polygon of H-transfer angle from NADPH (black) or C226 (red) to the C17 position of Pchlide. **D.** Frequency polygon of H-transfer distance from Cys226 (black) or NADPH (red) to the C18 position of Pchlide. **E.** Frequency polygon of H-transfer angle from NADPH (black) or C226 (red) to the C18 position of Pchlide. All MD simulations were run for 10 ns of protein-restrained equilibration and 100 ns unrestrained with three repeats and most stable run selected.

Further investigation of the catalytic mechanism was carried out using MD simulations on the C226A and Q248A variants. In our new structural model of the *T. elongatus* structural model of the POR ternary complex, Gln248 is located in the active site where it would be close enough to interact with the central Mg atom of Pchlide. MD simulations on the Q248A variant show that the Ala248 residue has moved away from the active site, possibly due to the loss of the adduct with the central Mg atom, which is likely to be important for Pchlide binding. Moreover, there is also a gap between the nicotinamide ring of NADPH and Cys226 that can now accommodate a water molecule, which may potentially cause changes to the Pchlide electronic structure that result in the increase in rate of the hydride and protein transfer steps in the Q248A variant (**Figure 9A**). The wild type simulations also highlight another, as yet unexplored, hypothetical route for ‘hydride’ transfer from NADPH via the Cys226 residue (**Figure 9B**). In this case the proposed hydrogen atom transfer, which follows an initial excited state electron transfer from NADPH to the C17-C18 double bond of Pchlide [10], could possibly proceed from Cys226 to the C18 position of Pchlide, as the distance and angle of transfer is close to optimal. This would be accompanied by a simultaneous hydrogen atom transfer from the NADPH to Cys226, although it should be noted that there is currently no experimental evidence to support or refute such a mechanism. However, it is evident from both the present and previous experimental data [9, 23] that any changes to this Cys residue affect both the hydride and proton transfer steps, so it is clearly important for active site chemistry.

**Fig. 9.**
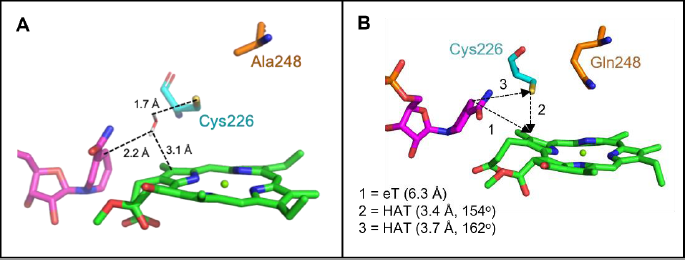
Potential implications of new structural model for POR catalysis. **A.** MD simulation of Q248A variant. Dotted lines represent potential hydrogen bonds (all < 5 Å) between NADPH, Cys226, Pchlide and a water molecule. **B.** MD simulation of wild-type POR, illustrating the potential for Cys226 to play a role in hydride transfer from NADPH to the C18 position of Pchlide. Putative direct electron transfer from NADPH to C18 (1), and simultaneous hydrogen atom transfer (HAT) routes from Cys226 to the C18 position of Pchlide (2) and from NADPH to Cys226 (3) are shown. All MD simulations were run for 10 ns of protein-restrained equilibration and 100 ns unrestrained with three repeats.

## CONCLUSIONS

As different structural models have previously been hypothesised for the ternary enzyme-substrate complex of POR based on cryo-EM and crystallography data [12, 14], we have now provided deeper insights into the validity of each model through a combination of computational and experimental approaches. The site-directed mutagenesis studies presented here are compatible with a structural conformation similar to the ‘closed’ structure of the cryo-EM model [14]. Although a conserved Tyr residue has been widely assumed to act as the putative proton donor in POR photocatalysis [8, 15], it is unlikely to perform such a role, but instead is crucial for Pchlide binding. The conserved Cys226 residue is located close to the C18 of the Pchlide substrate and is essential for active site chemistry. Our studies suggest it is likely to act as the proton donor, either directly or via a water-mediated network, and it also appears to play a key role in the hydride transfer step. Mutation of Cys226 causes a significant movement of the nicotinamide ring below the plane of Pchlide which results in the altered mechanism observed in Cys226 variants [9, 23]. An additional active site residue, Gln248, is likely to interact with the central Mg atom of Pchlide and has been shown to be important for binding and tuning the electronic properties of the Pchlide substrate for efficient photochemistry. Based on these findings a new ‘binding pose’ is presented for the *T. elongatus* POR ternary enzyme-substrate complex that is compatible for reaction chemistry. However, confirmation would ultimately require a crystal structure of the POR-Pchlide-NADPH ternary complex, which may also open up exciting new opportunities for time-resolved visualisation of the light-driven reaction chemistry.

## EXPERIMENTAL PROCEDURES

### Materials

All chemicals were obtained from Sigma-Aldrich apart from auto induction Luria-Bertani (LB) media and Tris-base, which were purchased from Formedium (Norfolk, UK), and 3-fluoro-L-tyrosine (monoF-L-tyrosine = mFY), which was purchased from Tokyo Chemicals Industry UK Ltd (Oxford, UK). 3,5-difluoro-L-tyrosine (diF-L-tyrosine = dFY) and 2,3,5trifluoro-L-tyrosine (triF-L-tyrosine = tFY) were synthesised by a modification of a previously described procedure [29]. For the synthesis of dFY ammonium acetate (16.6 g, 0.2 M) and sodium pyruvate (11.0 g, 0.1 M) were combined in 800 mL H_2_O, and the pH was adjusted to 8.2 with about 1 mL of 7 M NH_3_. The volume was increased to 1 L, then 13.7 mg pyridoxal phosphate and 0.35 mL of 2-mercaptoethanol were added. 2,6-difluorophenol (1.30 g, 10 mmol) was then added, and 1.0 mL (10 mg/mL) of tyrosine phenol lyase solution. 2,6-difluorophenol was added in 0.65 g portions each day for the next three days. The reaction mixture was then loaded on a column of Dowex-50 (H^+^) (5 x 40 cm) and the column washed with water until the eluate was neutral. The product was eluted with 200 mL 2 M NH_3_, followed by 200 mL water. The combined eluates were evaporated in vacuo, and the residue was crystallised from hot water to give 2.7 g (49.6 %) white granular product. NMR (**Figures S14 and S15**): ^1^H (D_2_O, 400.1 MHz), δ 6.79 (2H, d, J=8 Hz), 4.18 (1H, dd, J=5.6, 7.4 Hz), 3.12 (1H, dd, J=5.4, 14.8 Hz), 2.995 (1H, dd, J=7.6, 14.8 Hz); ^19^F (D_2_O, 376.5 MHz), δ -133.16 (d, J=8.5 Hz). For the synthesis of tFY the reaction was performed as above, using 2.25 g 2,3,6-trifluorophenol. Yield, 0.80 g (23%) light yellow solid. NMR (**Figures S16 and S17**): ^1^H (D_2_O, 400.1 MHz), δ 6.79 (2H, m), 4.18 (1H, t, J=6.9 Hz), 3.196 (1H, dd, J=5.7, 14.9 Hz), 3.058 (1H, dd, J=7.2, 15.0 Hz); ^19^F (D_2_O, 376.5 MHz), δ -138.19 (dt, J=8.2, 11.3 Hz), -145.318 (m), -156.0 (dq, J=2.6, 8.3, 12.0 Hz).

The POR gene from *Thermosynechococcus elongatus* was codon optimised for overexpression in *E. coli*, synthesised (GeneArt, ThermoFisher) and sub-cloned into pET21a with a C-terminal 6-His tag. The pEVOL_FNY plasmid, containing 2 copies of aaRS and the tRNA for incorporation of fluorinated Tyr analogues, were obtained from Prof. JoAnne Stubbe (Massachusetts Institute of Technology, USA). The Pchlide substrate was produced and purified as previously described [12].

### Site-directed mutagenesis of T. elongatus POR

All primers were designed using the online QuikChange Primer Design program and are shown in **Table S1**. PCR reactions were performed according to conditions in [12]. PCR products were purified (gel extraction; Qiagen, Hilden, Germany), incubated with Dpn I (New England Biolabs Inc., Ipswich, MA, USA) and transformed into *E. coli* NEB5α. Transformed cells were selected on (LB) agar plates (ampicillin 100 μg mL^−1^) and incubated overnight at 37 °C. Single colonies were selected, and grown in LB medium (ampicillin 50 μg mL^−1^) and plasmid purified (miniprep kit Qiagen, Hilden, Germany). Mutations were confirmed by DNA sequencing (Eurofins Genomics, Ebersberg, Germany).

### Expression and purification of T. elongatus POR proteins

Recombinant protein samples were prepared from *E. coli* BL21(DE3) transformed with the POR genes in pET21a and grown at 25 °C overnight in 2 L flasks with 500 mL auto induction LB medium containing 50 μg mL^−1^ ampicillin. For the fluorinated tyrosine variants, *E. coli* BL21 (DE3) cells were co-transformed with the POR gene (containing the amber stop codon for addition of the fluorinated tyrosine) in pET21a and a vector for the translational component system (pEVOL_FNY) and grown in 500 mL 2xYT medium containing 100 μg mL^−1^ ampicillin and 25 μg mL^−1^ chloramphenicol. The cultures were grown at 37°C until an OD600 of 0.3, at which point the 2 mM mFY, dFY or tFY was added. When the OD reached 0.5 expression was induced using 100 μM IPTG and 0.05 % arabinose and cultures grown at 25 °C overnight. After harvesting, cells were resuspended in 25 mm imidazole, 0.1 % (v/v) 2-mercaptoethanol, 150 mm NaCl, 20 mm HEPES, pH 7.5 and lysed by sonication (30 × 15 s). After centrifugation (20 000 g; 30 min; 4 °C) the supernatant was loaded onto a 5-mL HisTrap HF column and washed with different concentrations of imidazole in the washing buffer [0.1% (v/v) 2-mercaptoethanol, 150 mm NaCl, 20 mm HEPES, pH 7.5]. POR protein was eluted using elution buffer [250 mm imidazole, 0.1 % (v/v) 2-mercaptoethanol, 150 mm NaCl, 20 mm HEPES, pH 7.5] and concentrated in a Vivaspin concentrator (Generon Ltd, Slough, UK). Imidazole was removed by a PD10 desalting column (Biotech GmbH, Berlin, Germany), protein concentration was determined by absorbance at 280 nm (using an extinction coefficient of 35.41 mm−1·cm−1) and purity shown by SDS/PAGE. Protein was desalted into alternative buffer for pH dependence measurements containing 50 mM MES, 25mM Tris, 25 mM ethanolamine, 100 mM NaCl.

### Steady-state kinetic assays

All samples for steady-state kinetic measurements were prepared under low-intensity green light and the assays carried out as in [19]. Substrate or product concentrations in the reaction buffer [100-500 μM NADPH, 1–30 μM Pchlide, 0.1% Triton X-100, 0.1% (v/v) 2-mercaptoethanol, 150 mM NaCl, 20 mM HEPES, pH 7.5] were determined using the following extinction coefficients in aqueous solution: NADPH, 6.22 mM^−1^·cm^−1^ at 340 nm; Pchlide, 24.95 mM^−1^·cm^−1^ at 630 nm; and Chlide, 69.95 mM^−1^·cm^−1^ at 670 nm. Steady-state activity measurements were carried out at 40 °C using a Cary 50 spectrophotometer (Varian Inc) to measure the initial rates of Chlide formation of 0.1-0.2 µM POR over a range of Pchlide concentrations. A 625 nm high power LED (Thorlabs Inc., Newton, NJ, USA) was used to provide continuous illumination (∼100 µmol photons·m^−2^·s^−1^) during the reaction assay. The apparent *K*_m_ and *V*_max_ values were obtained by fitting the initial rates of Chlide synthesis against the concentration of substrate (Pchlide or NADPH) to the Michaelis–Menten equation. The apparent *k*_cat_ values were calculated based on the reaction with close to saturated NADPH (200 µM) and varying Pchlide (up to 30 µM).

### Pchlide binding assays

The red spectral shift in the Pchilde absorption to 642 nm on formation of the POR ternary complex was used to follow Pchlide binding [30]. Binding assays comprised 0.5 μM Pchlide, 100 μM NADPH and 0.5–200 μM POR. Apparent *K*_d_ values were obtained by fitting the absorbance ratio changes (Δ*A* = *A*_642 nm_/*A*_630 nm_) against the concentration of POR using the equation (*A*_o_ is the initial ratio of the absorbance at *A*_642 nm_/*A*_630 nm_):

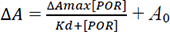

### Cryogenic absorbance spectroscopy

Low-temperature absorbance spectra were measured in reaction buffer with 44% glycerol, 20% sucrose and 0.1% Genapol in a Cary 50 spectrophotometer (Varian Inc., Palo Alto, CA, USA). 1 mL of samples containing 10 μM Pchlide, 250 μM NADPH and 50 μM POR were incubated at the corresponding temperature using an Optistat DN liquid nitrogen cryostat (Oxford Instruments Inc., Abingdon, UK). The POR reaction was initiated by the same LED light used in the activity measurements. For each temperature point, the sample was illuminated for 10 min, or for the dark steps, the sample was illuminated at 190 K for 10 min, and then incubated in the dark at the corresponding temperature for 10 min. All absorbance spectra were recorded at 77 K by using a Cary 50 spectrophotometer (Agilent Technologies, Santa Clara, CA, USA).

### Laser flash photolysis

Laser photoexcitation experiments were carried out at 25°C using an Edinburgh Instruments LP980 transient absorption spectrometer. Samples of the POR ternary complex (experimental conditions are provided in the figure legends) excited at 450 nm (∼15 mJ), using the optical parametric oscillator of a Q-switched Nd-YAG laser (NT342B, EKSPLA) in a cuvette of 1 cm pathlength. Rate constants and amplitudes were calculated from the average of five time-dependent absorption measurements by fitting to a single exponential function in Origin Pro 9.1 software. Relative quantum yields were calculated by measuring the amount of Chlide produced per laser pulse and normalised relative to the amount of bound Pchlide.

### Molecular Dynamics

All MD simulations were carried out in Gromacs 5.0.432 using the Amber14 force field with the system in a solvation box of at least 12 Å with counter-ions generated in AmberTools, retaining crystallographic waters, totalling 48,561 atoms for the wild type ternary enzyme-substrate complex. The following parameters were used: constant pressure (1 bar), 10 Å van der Waals and electrostatic cut-offs, particle mesh Ewald for long-range electrostatics, LINCS bond restraints and periodic boundary conditions and a 2 fs time step. Charge parameterisation and bonding parameters for Pchlide and NADPH are from Zhang et al [12].

### Binding energy calculations

Binding energy calculations were carried out by calculating Gromacs free energy using the slow-growth approach which gradually decouples the substrate from the system [30]. A coupling parameter, λ, was used to indicate the level of change in a system between two states A (coupled) and B (fully decoupled), ΔG_AB_. Simulations were conducted using a range of λ values, and results were plotted as ∂H/∂λ, from which ΔG_AB_ was derived.

## Abbreviations

POR: protochlorophyllide oxidoreductase
Pchlide: protochlorophyllide
Chlide: chlorophyllide
cryo-EM: cryo-electron microscopy

## Notes

### Competing Interest Statement

The authors have declared no competing interest.

