## supplemental information for "Mechanistic implications of the ternary complex structural models for the photoenzyme protochlorophyllide oxidoreductase"

### Supplementary Figures

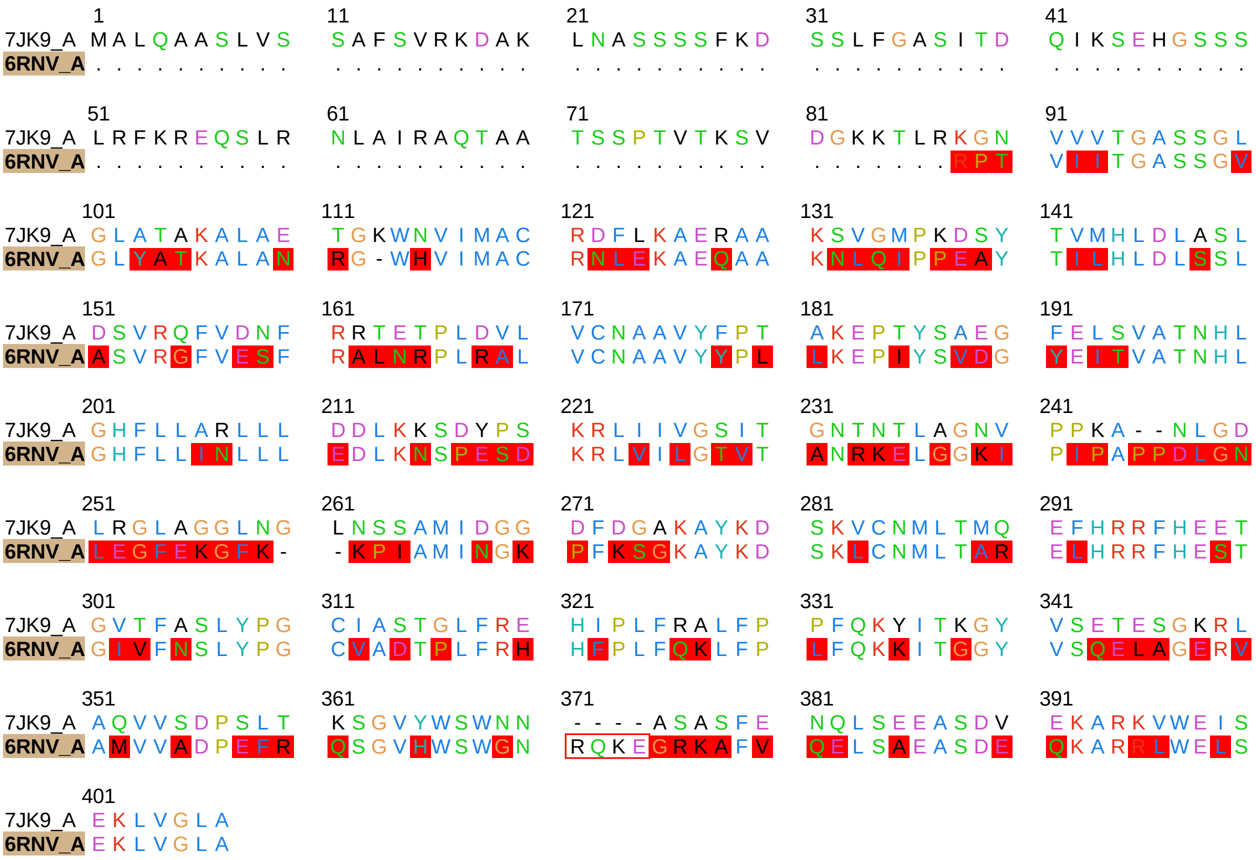


**Figure S1. Sequence alignment of *A. thaliana* POR (7JK9) [1] with *T. elongatus* (6RNV) [2] using Chimera multi-aligner**. 57 % sequence match; non-matching amino acids highlighted in red. Numbering is based on the sequence of the *A. thaliana* POR.

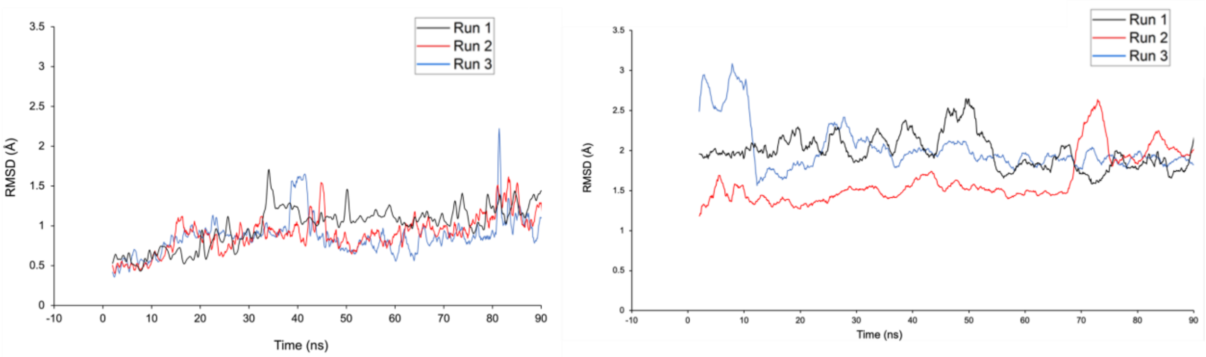


**Figure S2. Initial MD simulations showing stability of proposed model of *T. elongatus* [12] and *A. thaliana* [14] POR-NADPH-Pchlide.** MD simulations were run for 10 ns of protein-restrained equilibration and 100 ns unrestrained. RMSD of Pchlide was measured during MD simulations using *T. elongatus* [12] (**left**) and *A. thaliana* [14] **(right)** relative to its starting position for all atoms except hydrogen; Run 1 = black, Run 2 = red, Run 3 = blue. Data is plotted to a moving average (interval 0.5 ns).

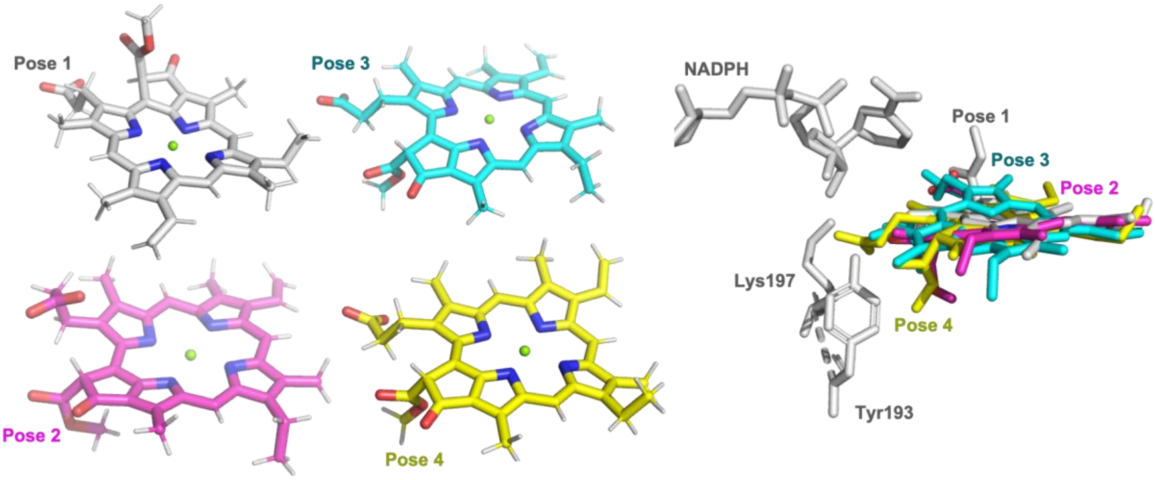


**Figure S3. Investigation of the different potential binding poses of Pchlide in the active site of POR.** The three highest scoring Pchlide docking poses from AutoDock Vina (pose 1-3) and a Pchlide binding pose based on the model proposed by Nguyen et al [14] (pose 4) are compared individually (left) and overlaid in in the active site (right) with NADPH, Lys197 and Tyr193 from the pose 1 model (i.e. the same conformation suggested by Zhang et al [12]).

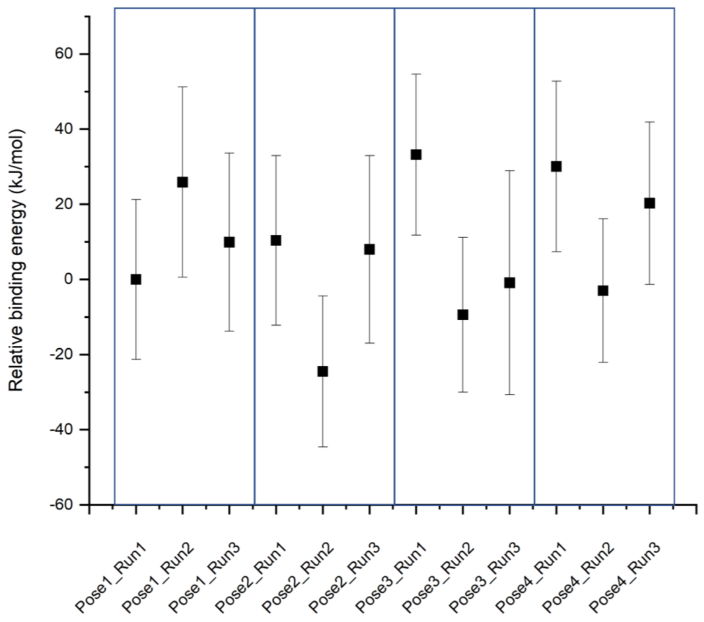


**Figure S4. Free energy compared across 4 different poses of Pchlide in POR active site.** Data are shown relative to Pose1_Run1. MD simulations were run for 10 ns of protein-restrained equilibration and 110 ns unrestrained. The energies were calculated using Gromacs slow-growth method [3].

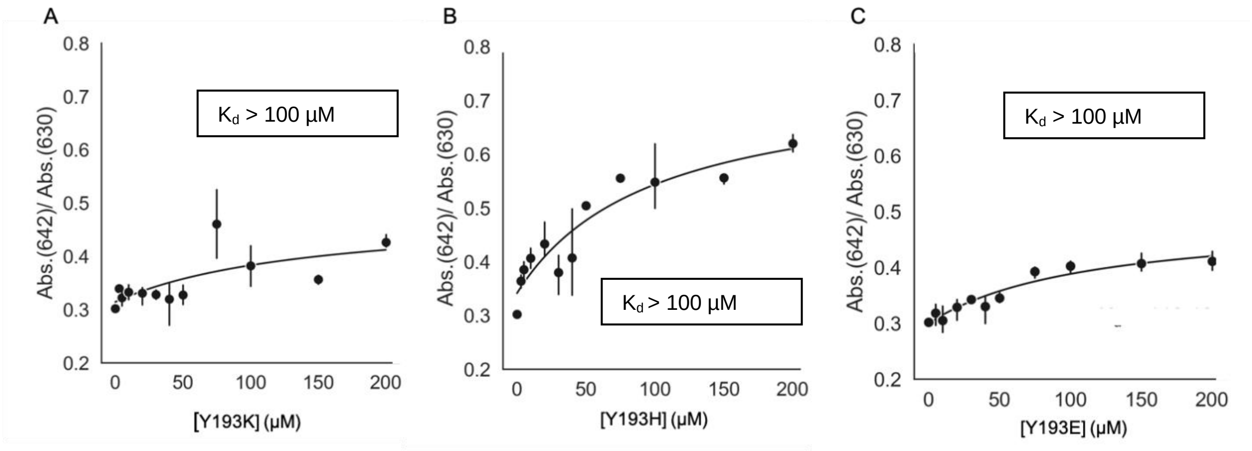


**Figure S5. Pchlide binding analysis of Y193 POR variants**. Pchlide binding titrations for Tyr193 variants of *T. elongatus* POR, measured by plotting the absorbance ratio of bound Pchlide (642 nm) against free Pchlide (630 nm) upon increasing concentrations of enzyme. Reaction conditions: 0.5 µM Pchlide, 0.1 % Triton, 1 µM 2-mercaptoethanoland 100 µM NADPH. **A** Y193K variant, **B** Y193H variant, **C** Y193E variant. Error bars depict standard deviation of the mean (n=3).

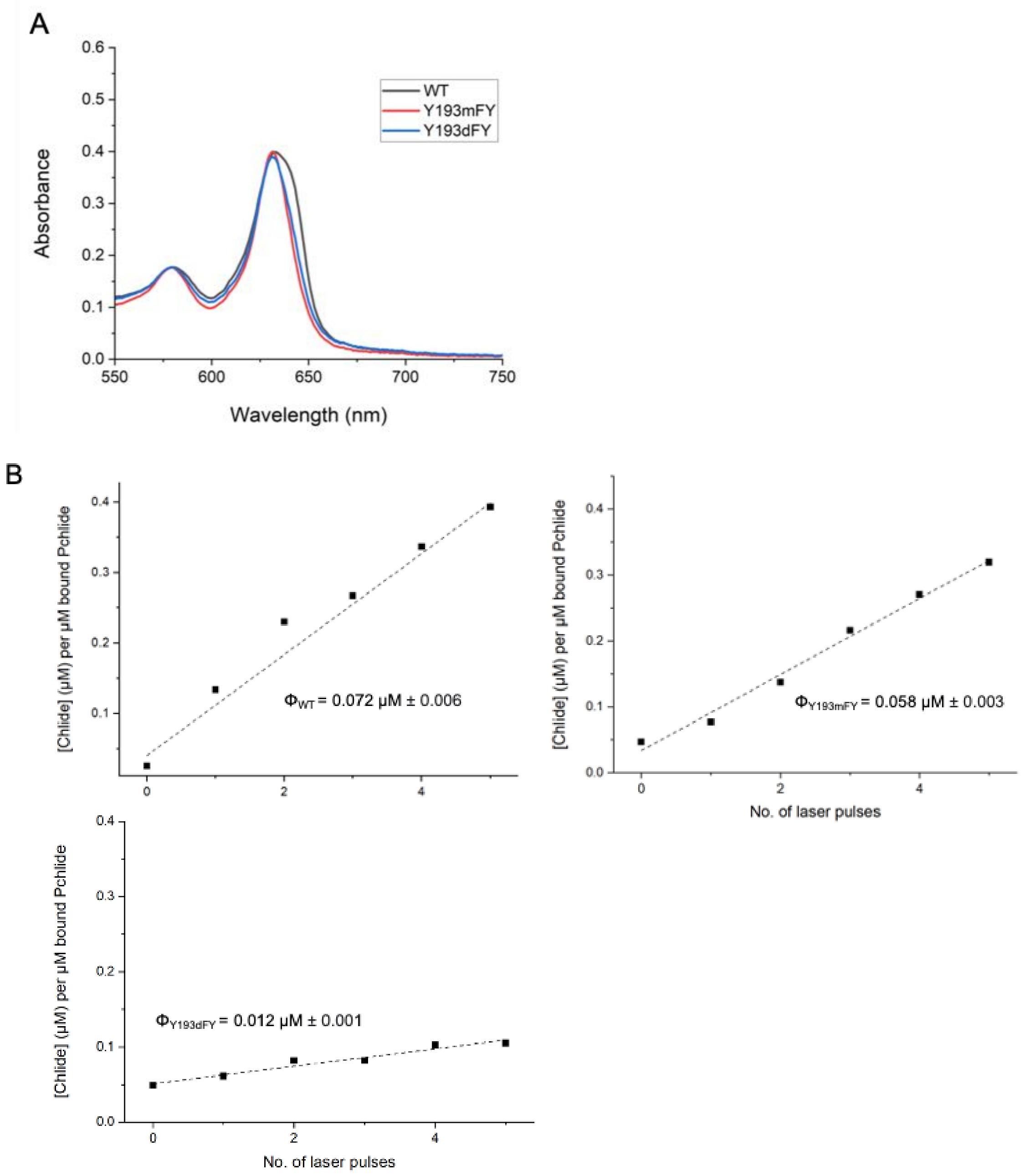


**Figure S6**. **Relative quantum yield for wild type and fluorinated POR variants. A** ‘Bound ternary complex’ absorbance spectrum of wild-type (black), mono-fluorinated Tyr193 POR (Y193mFY, red) and di-fluorinated Tyr193 POR (Y193dFY, blue) at enzyme concentrations of 100 µM, 200 µM and 400 µM, respectively. The relative percentage of bound Pchlide compared to wild type was calculated to be 100%, 70% and 70%, respectively. **B** The concentration of Chlide produced after each laser pulse was plotted (normalised using bound Pchlide concentration) and fitted to a straight line with the slope used as an estimate of the relative quantum yield for the POR reaction. Reaction conditions: 15 µM Pchlide, % Triton, 1 µM 2-mercaptoethanol, 250 µM NADPH, 100 µM WT/ 200 µM Y193mFY/ 400 µM Y193dFY.

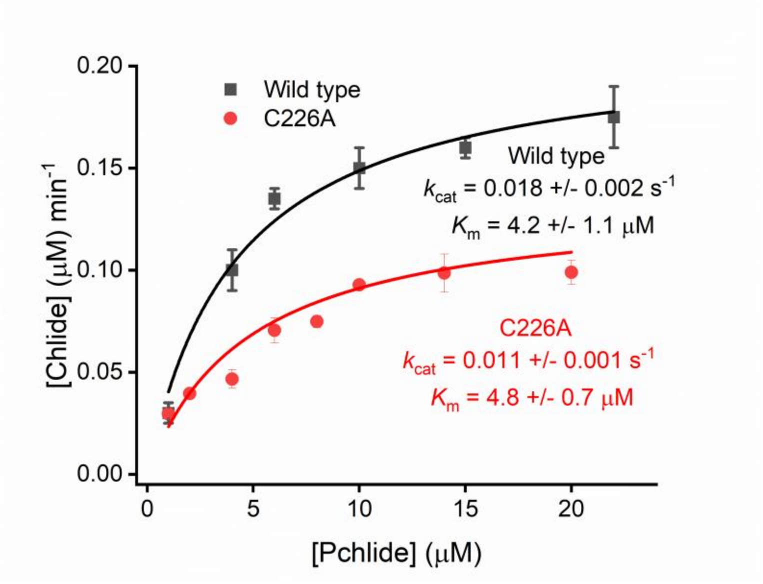


**Figure S7. Steady-state kinetics analysis for C226A POR variant compared to wild type POR.** Michaelis Menten plots showing the rate of Chlide formation (by measuring absorbance increase at 670 nm) for C226A POR variant (red) and wild type POR (black) measured over a range of Pchlide concentrations. Reaction conditions: 0.2 µM POR, varying concentrations of Pchlide, 0.1 % Triton, 1 µM 2-mercaptoethanoland 100 µM NADPH.

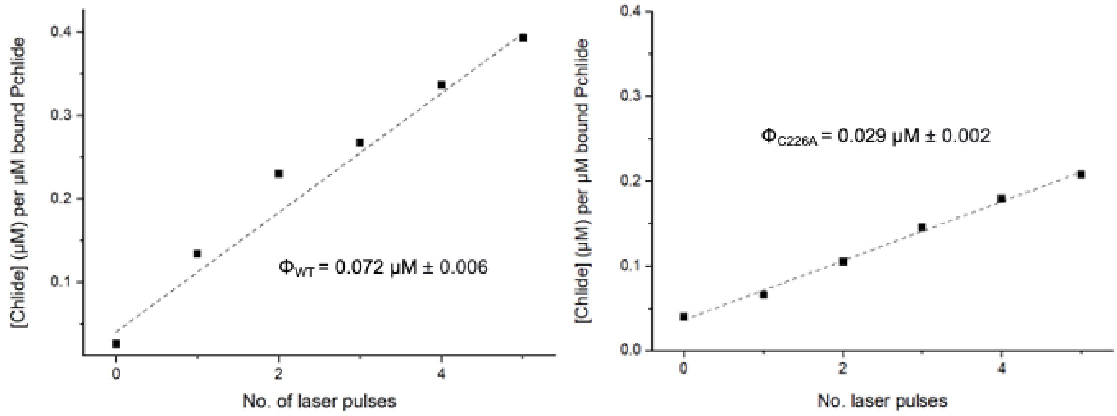


**Figure S8. Relative quantum yield for wild type and C226A variant.** Reaction conditions: 15 µM Pchlide, 100 µM wild type, 0.1 % Triton, 1 µM 2-mercaptoethanoland 250 µM NADPH (left), 30 µM Pchlide, 200 µM C226A, 0.1 % Triton, 1 µM 2-mercaptoethanoland 500 µM NADPH (right). The Chlide concentration after each laser pulse was plotted (normalised using bound Pchlide concentration) and fitted to a straight line with the slope used as an estimate of the relative quantum yield for the POR reaction.

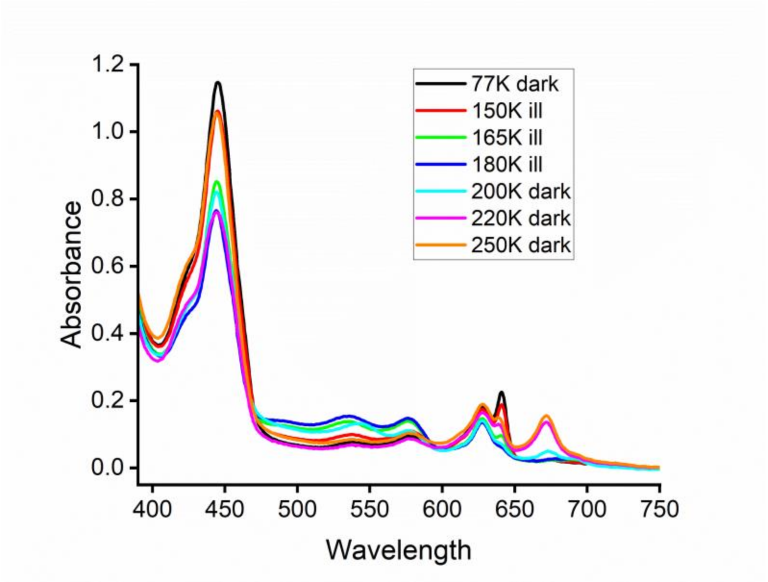


**Figure S9. Cryogenic absorbance measurements of POR C226A variant.** Samples contained 100 µM C226A, 20 µM Pchlide, 0.1 % Triton, 1 µM 2-mercaptoethanoland 250 µM NADPH in 20 % sucrose, 44 % glycerol, 150 mm NaCl and 20 mm HEPES pH 7.5. The absorbance spectra of the C226A-Pchlide-NADPH ternary complex were measured at 77 K after illumination for 10 min at various temperatures. For the ensuing light-independent or dark step absorbance spectra of the C226A-Pchlide-NADPH ternary complex were measured at 77 K after illumination for 10 min at 180 K followed by incubation in the dark at various temperatures.

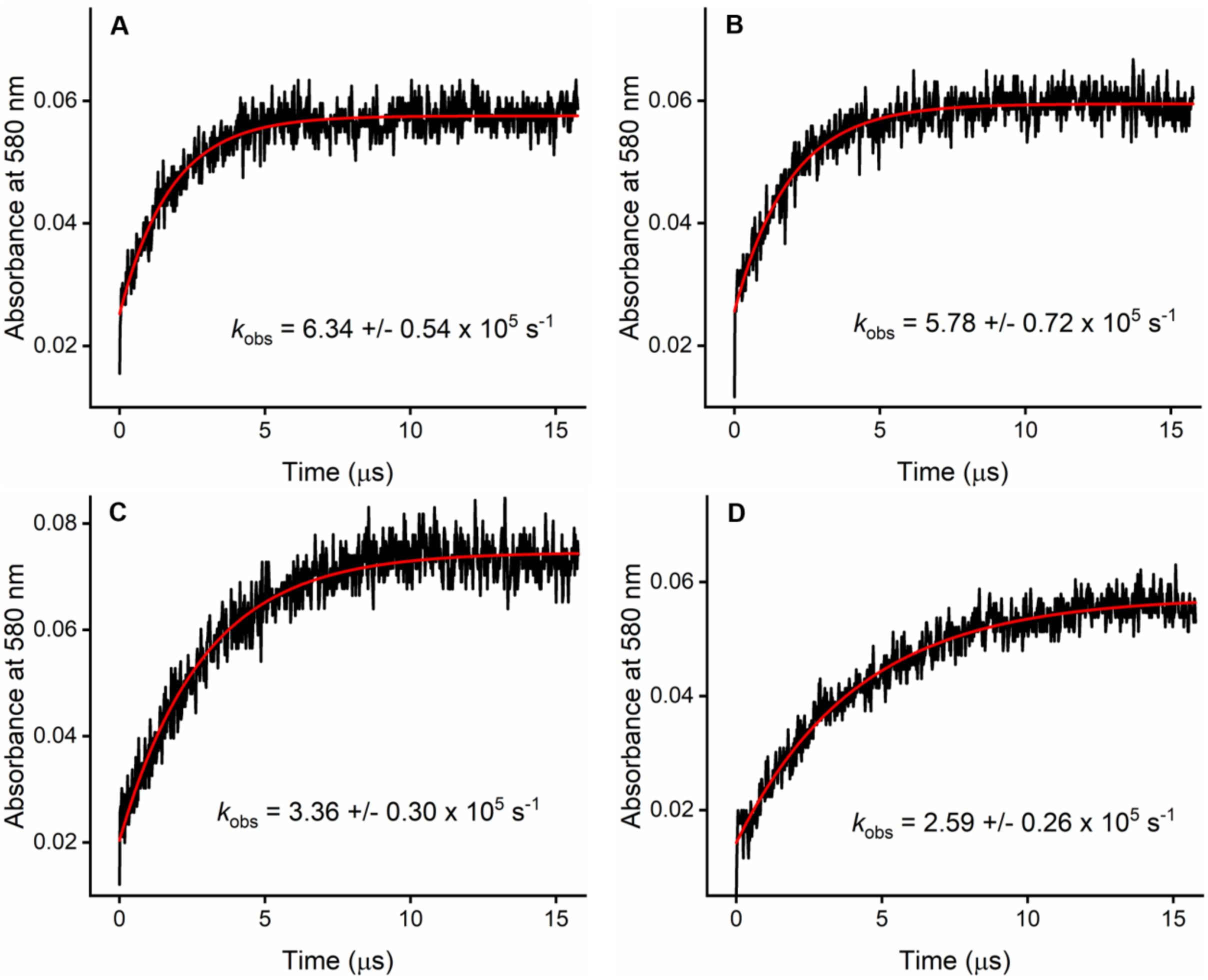


**Figure S10. Laser photoexcitation measurements illustrating the rate of hydride transfer for C226A POR variant under different deuteration conditions.** Absorbance transients at 580 nm after excitation with a laser pulse at 450 nm (15 mJ) are shown for C226A ternary complex samples together with: **A**. NADPH in $H_2O$ buffer, **B**. NADPH in $D_2O$ buffer, **C**. NADPD in $H_2O$ buffer and **D**. NADPD in $D_2O$ buffer. Reaction conditions: 100 µM C226A, 20 µM Pchlide, 0.1 % Triton, 1 µM 2-mercaptoethanoland 250 µM NADPH/D.

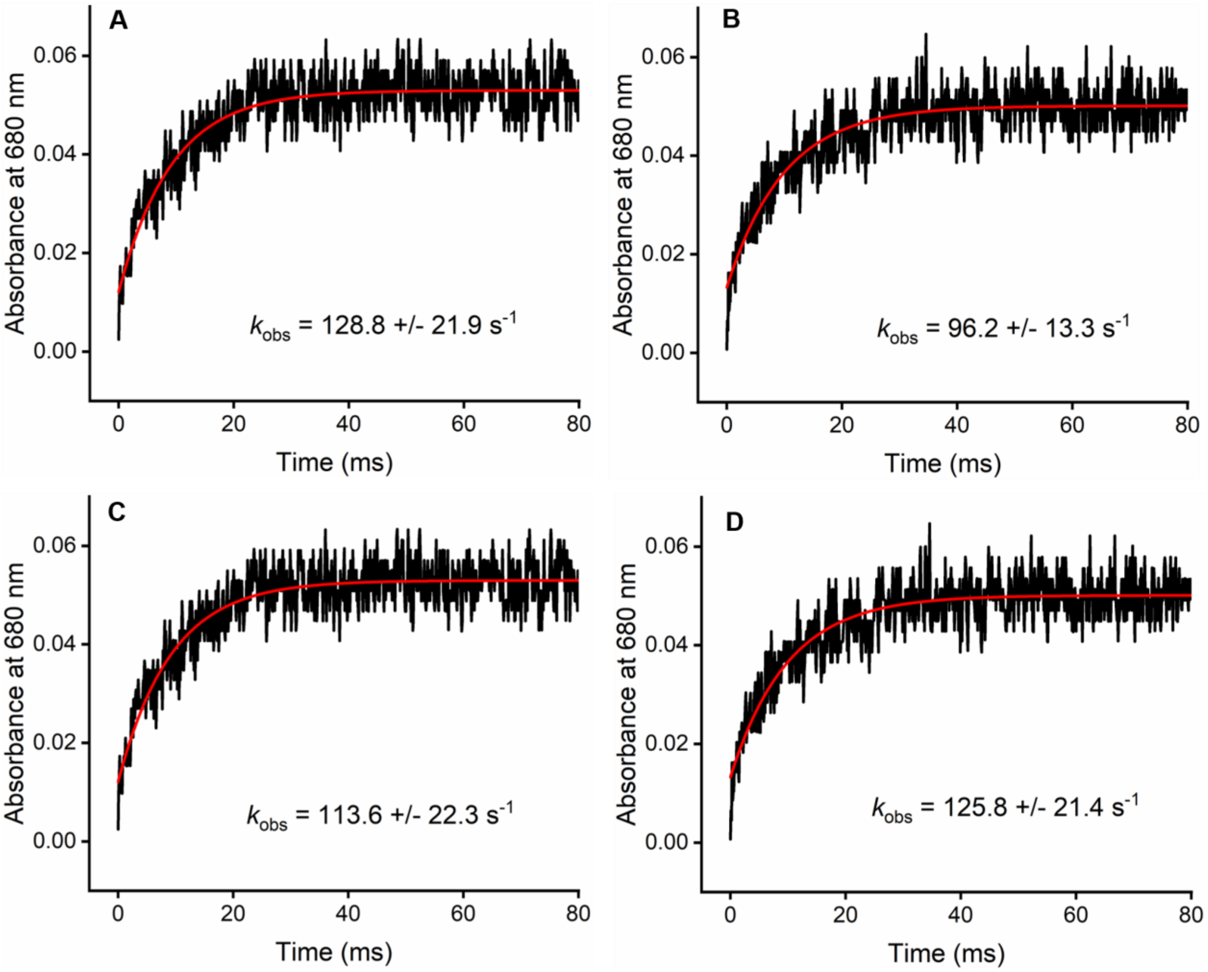


**Figure S11. Laser photoexcitation measurements illustrating the rate of proton transfer for C226A POR variant under different deuteration conditions.** Kinetic transients at 680 nm after excitation with a laser pulse at 450 nm (15 mJ) are shown for C226A ternary complex samples together with: **A**. NADPH in $H_2O$ buffer, **B**. NADPH in $D_2O$ buffer, **C**. NADPD in $H_2O$ buffer and **D**. NADPD in $D_2O$ buffer. Reaction conditions: 100 µM C226A, 10 µM Pchlide, 0.1 % Triton, 1 µM 2-mercaptoethanoland 250 µM NADPH/D.

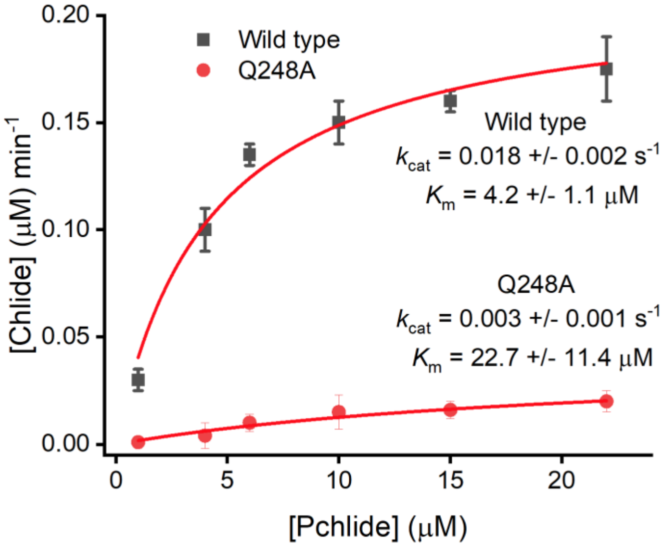


**Figure S12. Steady-state kinetics analysis for Q248A POR variant compared to wild type POR.** Michaelis Menten plots showing the rate of Chlide formation (by measuring absorbance increase at 670 nm) for Q248A POR variant (red) and wild type POR (black) measured over a range of Pchlide concentrations. Reaction conditions: 0.2 µM POR, varying concentrations of Pchlide, 0.1 % Triton, 1 µM 2-mercaptoethanoland 100 µM NADPH.

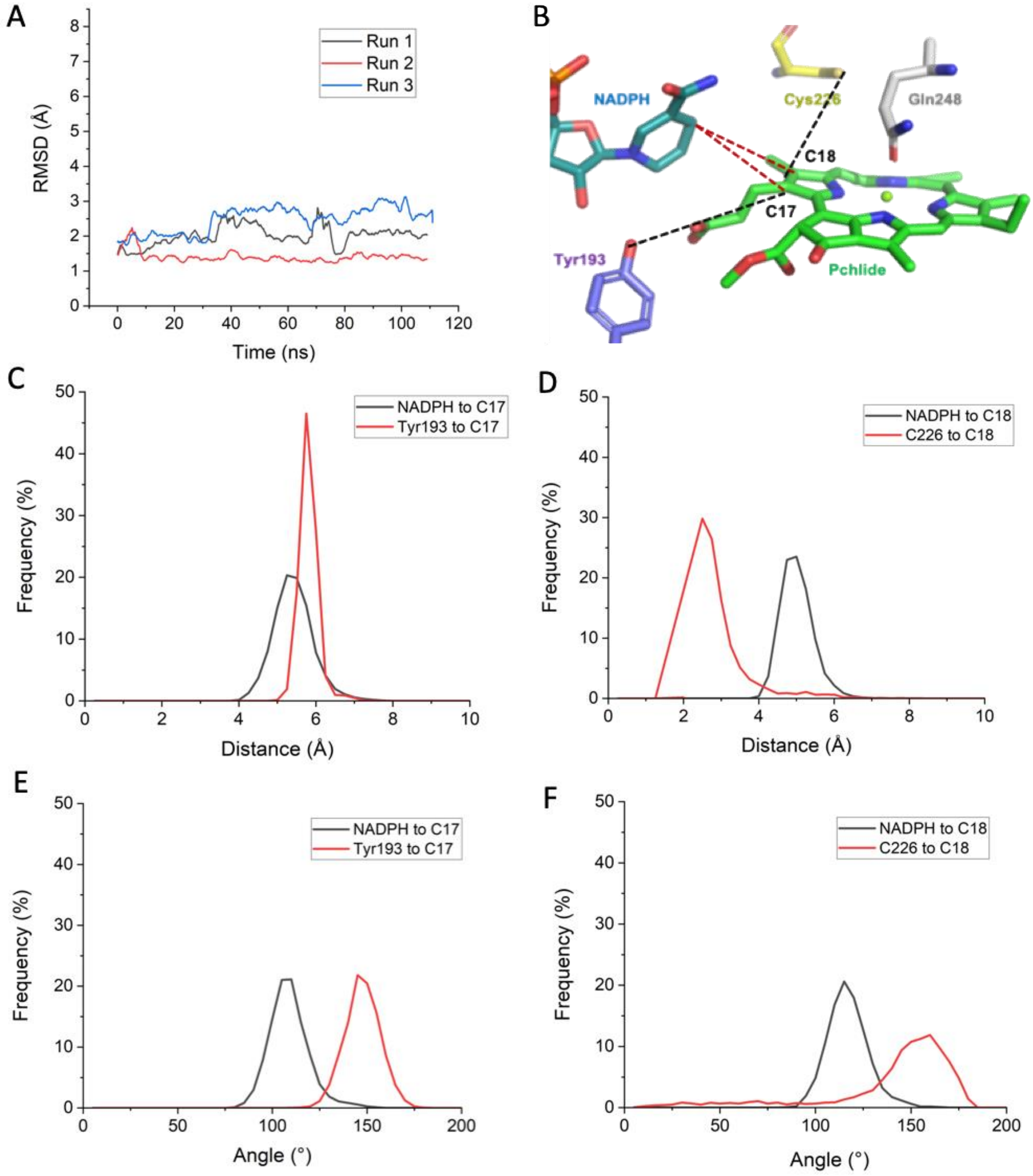


**Figure S13. MD analysis of the new structural model of the *T. elongatus* POR-Pchlide-NADPH ternary complex**. MD simulations were run for 10 ns of protein-restrained equilibration and 100 ns unrestrained **A.** RMSD of Pchlide relative to its starting position for all atoms except hydrogen; Run 1 (black), Run 2 (red), Run 3 (blue). Data is plotted to a moving average (interval 2 ns). **B.** Single-linkage cluster from most stable run (Run 2) demonstrating the key residues; Cys226, Gln248 and Tyr193. Potential hydride transfer from NADPH to the C17 or C18 positions of Pchlide is shown in red and potential proton transfer from either Tyr193 to the C17 position of Pchlide or Cys226 to the C18 position of Pchlide is shown in black. **C.** Transfer distance to the C17 position of Pchlide from Tyr193 (black) or NADPH (red). **D.** Transfer distance to the C18 position of Pchlide from Cys226 (black) or NADPH (red). **E** Angle of transfer to the C17 position of Pchlide from NADPH (black) or C226 (red). **F.** Angle of transfer to the C18 position of Pchlide from NADPH (black) or C226 (red).

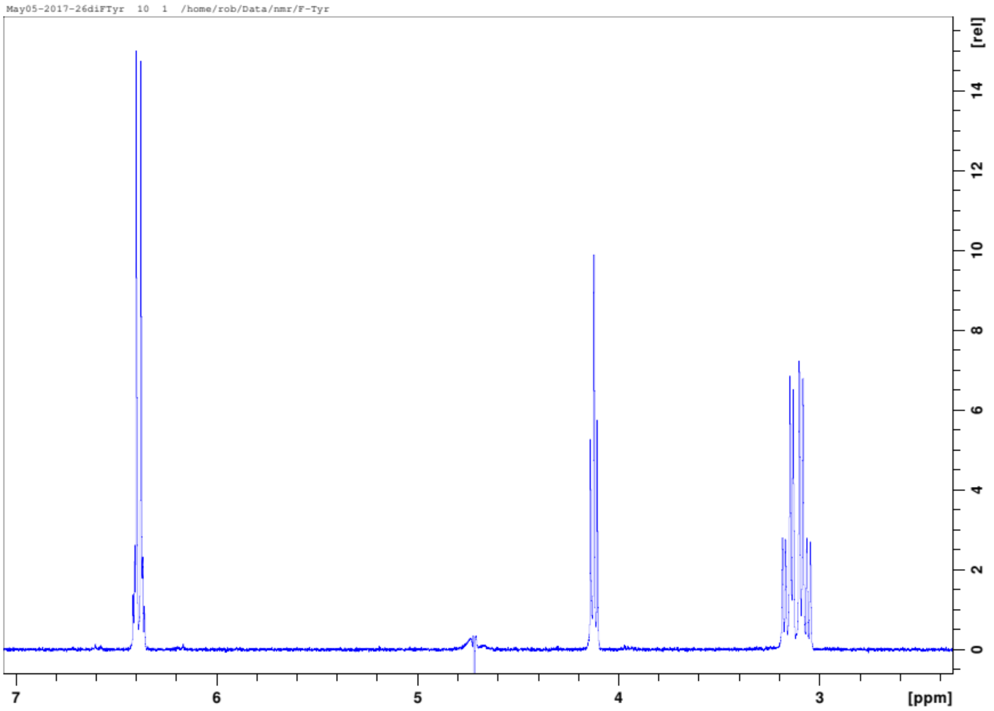


**Figure S14. $^1$H-NMR spectrum of 3,5-difluoro-L-tyrosine.**

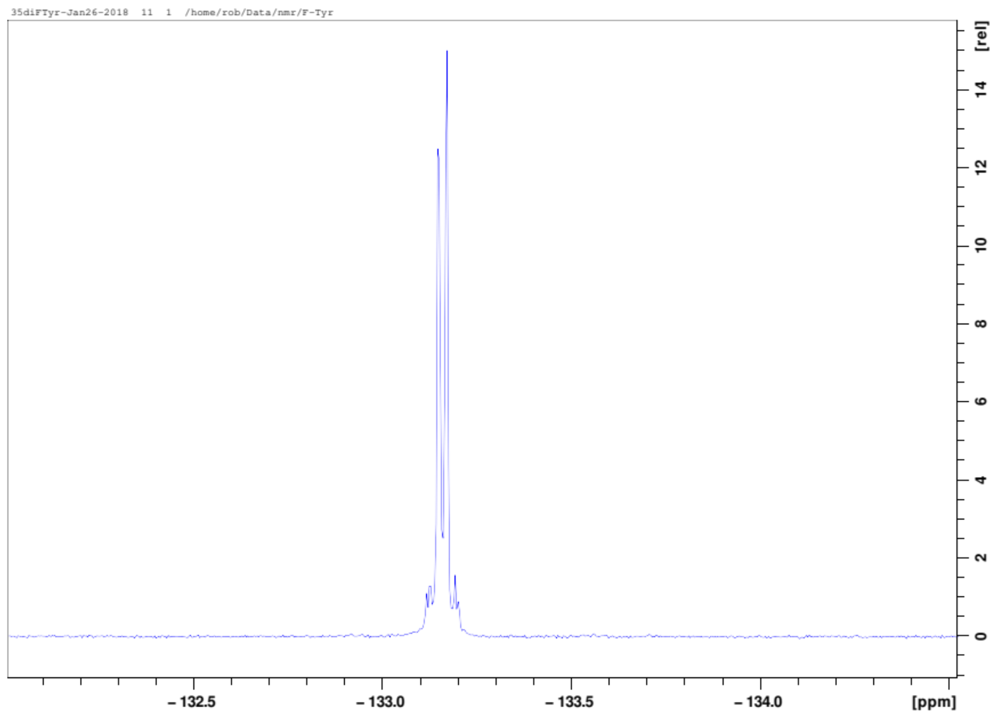


**Figure S15. $^{19}$F-NMR spectrum of 3,5-difluoro-L-tyrosine.**

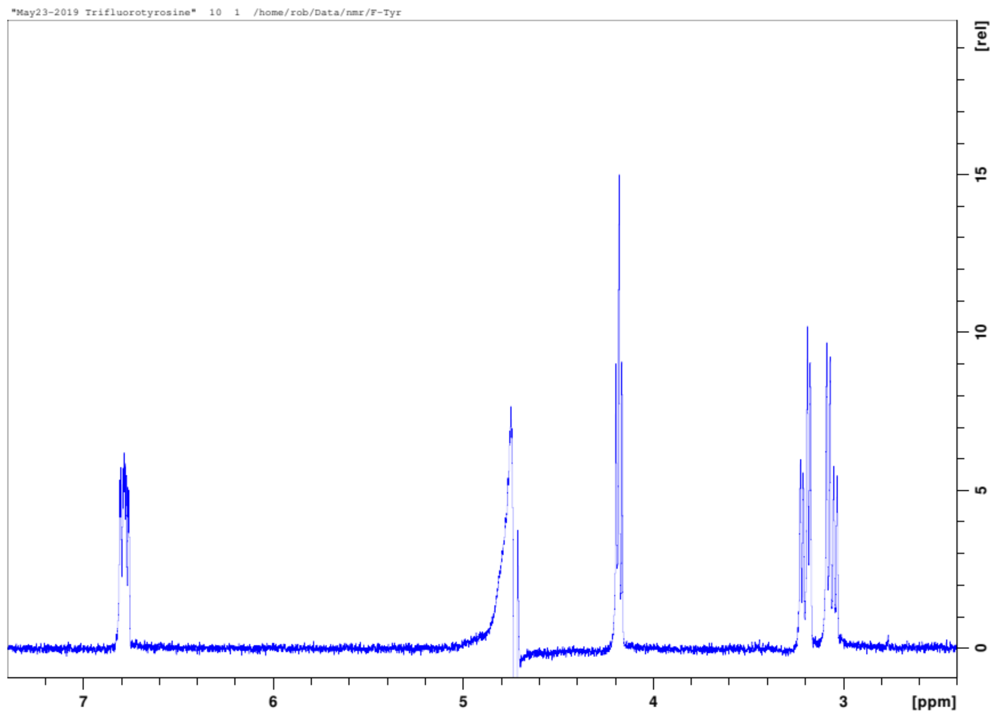


**Figure S16.** **$^{1}$H-NMR spectrum of 2,3,5-trifluoro-L-tyrosine.**

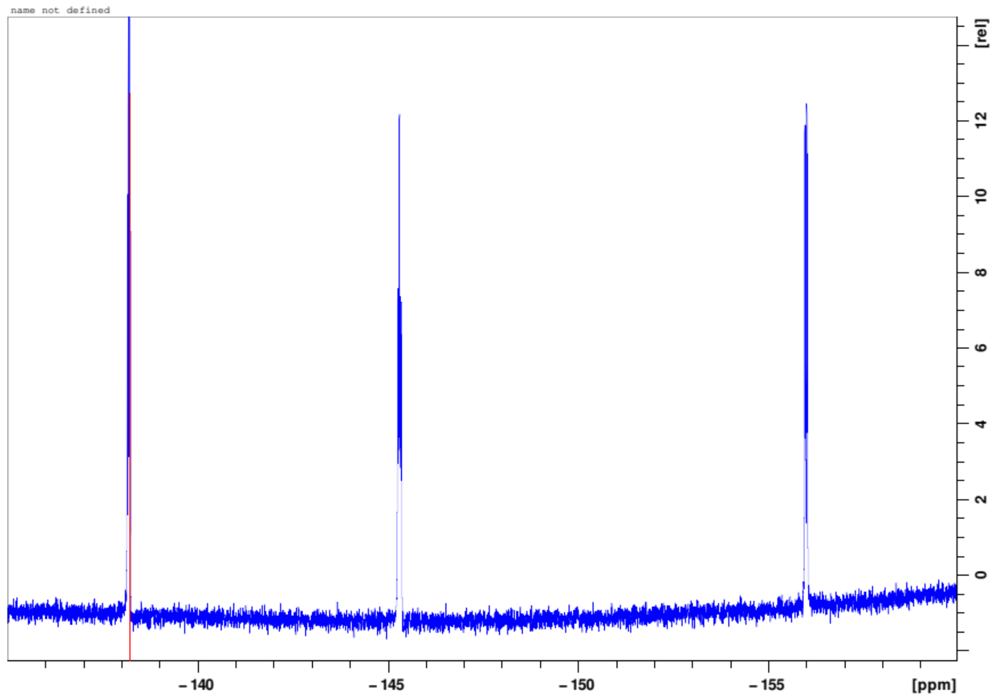


**Figure S17.** $^{19}$**F-NMR spectrum of 2,3,5-trifluoro-L-tyrosine.**

**Table S1.** Primers used for *T. elongatus* POR variants. Primer Sequence 5' to 3'

| POR variant | Primer sequence |
|---|---|
| Y193H | 5'-CACAGCTTGCTATCTTTATGGGCTTTGCCGCTTTTAAAC-3'<br>5'-GTTTAAAAGCGGCAAAGCCCATAAAGATAGCAAGCTGTG-3' |
| Y193E | 5'-ATTACACAGCTTGCTATCTTTCTCGGCTTTGCCGCTTTTAAACGG-3'<br>5'-CCGTTTAAAAGCGGCAAAGCCGAGAAAGATAGCAAGCTGTGTAAT-3' |
| Y193K | 5'-ATTACACAGCTTGCTATCTTTCTTGGCTTTGCCGCTTTTAAACGG-3'<br>5'-CCGTTTAAAAGCGGCAAAGCCAAGAAAGATAGCAAGCTGTGTAAT-3' |
| Y193_STOP | 5'-ATTACACAGCTTGCTATCTTTCTAGGCTTTGCCGCTTTTAAAC-3'<br>5'-GTTTAAAAGCGGCAAAGCCTAGAAAGATAGCAAGCTGTGTAAT-3' |
| C226A | 5'-GCGGTGTATCTGCAACAGCACCCGGATACAGGCTATTAA-3'<br>5'-TTAATAGCCTGTATCCGGGTGCTGTTGCAGATACACCGC-3' |
| G248A | 5'-CATAACCACCTGTAATCTTTTTTGCGAACAGCGGAAACAGTTTCTGAA-3'<br>5'-TTCAGAAACTGTTTCCGCTGTTCGCAAAAAAGATTACAGGTGTTATG-3' |
